# Structural Rewiring of IL-7R Dimerization by an Oncogenic Transmembrane Mutation Can Be Reversed by Rational Design

**DOI:** 10.64898/2026.02.17.706319

**Authors:** Qian Wang, Min Chen, Asma Lasram, Saana Vihuri, Angela Z. Chou, Weixin Bian, Zhiming Dai, Outi Haapanen, Giray Enkavi, Christoph Pollmann, Ilpo Vattulainen, Tiantian Cai, Jacob Piehler, James J. Chou

## Abstract

Mutations within the transmembrane domains (TMDs) of single-pass transmembrane receptors often cause aberrant, ligand-independent receptor signaling associated with diverse malignancies, but their mechanism of action remain largely unknown. These TMD mutations are generally not targetable as they are buried in membrane. Here, we determined the mechanism of a gain-of-function (GOF) TMD mutation of interleukin-7 receptor (IL-7R) associated with T-cell acute lymphoblastic leukemia, and addressed the possibility of directly targeting the TMD mutation by using rationally designed transmembrane helices to restore order to uncontrolled signaling. We find that the GOF mutation of IL-7R severely shifts the TMD homodimerization interface, causing the receptor to homodimerize in a geometry that activates downstream signaling independent of ligand. Designed transmembrane helices that interfere with the new interface, delivered with mRNA technology, selectively block ligand-independent but not ligand-dependent signaling. Our study provides a conceptual framework for understanding and repairing disease-causing TMD mutations of single-pass cytokine receptors.

## INTRODUCTION

Many single-pass cytokine and growth factor receptors harbor mutations in the transmembrane domain (TMD) that cause ligand-independent receptor activation and diseases. These mutations are often not cysteine substitutions that could force receptor oligomerization via disulfide formation. Hence, their effect on signaling should be attributed to changes in conformation and/or oligomerization. Delineating the disease mechanism of these mutations necessitates a deeper understanding of receptor mechanism beyond ligand-induced conformational changes, which may provide previously unrecognized opportunities for therapeutic interventions.

The first report of a single-pass transmembrane (TM) receptor harboring a major oncogenic TMD mutation was on the neu/p185 receptor (rat equivalent of the human erb-b2 receptor tyrosine kinase HER2), of which the residue V664 in the TMD is mutated to Glu^1^. Analogous oncogenic TMD mutation V659E in human HER2 has also been identified to be an oncogenic driver of lung adenocarcinomas and is associated with poor prognosis in non-small lung carcinoma patients^2^. Unexpectedly, these charge-introducing mutations within the membrane, which are generally perceived as disruptive, can cause constitutive activation of neu and HER2, respectively^1,2^. Although the precise mechanism of the TMD mutations remain elusive, the findings suggested that the receptor TMD is not merely a membrane anchor but plays an active role in receptor dimerization and activation^2^. Selective inhibition of TMD mutation-induced oncogenic signaling remains an unsolved problem.

In addition to receptor tyrosine kinases, oncogenic non-cysteine mutations in the TMD have been identified in several class I cytokine receptors. A S505N mutation in the thrombopoietin receptor (TpoR) results in constitutive TpoR activation and essential thrombocythemia^3,4^. This mutation is believed to promote a mode of TpoR TMD dimerization that activates downstream signaling independent of thrombopoietin^5^. TMD mutations leading to ligand-independent signaling have also been reported for members of the common γ-chain (γc; also known as IL-2Rγ) family of interleukin receptors. For example, a recurrent hotspot mutation I242N in the IL-4R TMD was found in patients with primary mediastinal large B-cell lymphoma and can lead to JAK/STAT activation that does not depend on IL-4^6^.

More importantly, the TMD of IL-7R has emerged as a hotspot of oncogenic mutations involving both cysteine and cysteine-less mutations and insertions accounting for about 10% of IL-7R mutations associated with T-cell acute lymphoblastic leukemias (ALL)^7–11^. IL-7R is also a member of the γc family of cytokine receptors that plays important roles in the development, proliferation, and homeostasis of T and B cells^12–14^. Under normal circumstances, IL-7R needs to specifically heterodimerize with γc upon IL-7 binding to allow activation of downstream STAT5 phosphorylation via the Janus family tyrosine kinases JAK1 and JAK3^14–17^. A previous structural study revealed that the TMDs of IL-7R and γc can specifically interact in the membrane with a knob-into-hole mode of association and this interaction synergizes with ligand-induced heterodimerization of IL-7R and γc in activating downstream signaling^18^. A transmembrane mutation of IL-7R (V253G) found in some patients with ALL, however, is able to activate IL-7R without the ligand^9,10^, possibly by directly promoting homodimerization of the intracellular JAK1 and trans-activation^19,20^.

In this study, we sought to uncover the structural and mechanistic basis by which the transmembrane mutation V253G causes ligand-independent IL-7R signaling, and explore the possibilities of interfering with such oncogenic signaling via rationally designed transmembrane helices. To this end, we combined the use of bicelle NMR technology, single-molecule FRET, bacterial two-hybrid assays, functional mutagenesis, as well as molecular dynamics (MD) simulation to investigate the coupling between the TMD structure and intracellular JAK1 activation exploited by the disease mutation to activate the receptor without ligand. We find that the V253G mutation can completely alter the intrinsic dimerizing property of the TMD, resulting in a new type of TMD homodimerization that synergizes with auto cross activation of JAK1s. By means of a transmembrane peptide antagonist, we achieved selective inhibition of oncogenic homomeric signaling of IL-7R V253G while not interfering with the ligand-dependent receptor activation via γc.

## RESULTS

### The V253G mutation caused TMD homotypic association interface to rotate by ∼170°

As mentioned above, the V253G mutation in the IL-7R TMD induces constitutive STAT5 phosphorylation in BaF3 cells^9^ (**Fig. S1A**). To independently test this result, WT or V253G IL-7R was expressed in BaF3 cells with or without γc, and STAT5 and JAK1 phosphorylation were measured as an indicator of receptor activation. Cell surface expression of WT and mutant IL-7Rs were very similar except for modestly higher level of the mutant when co-expressed with γc (**Fig. S1B-E)**. Co-expression of γc and WT IL-7R showed ∼5 times stronger p-STAT5 and ∼3 times stronger p-JAK1 upon IL-7 treatment (**Fig. S1F-G**), whereas V253G exhibited strong p-STAT5 and p-JAK1 signals in the absence of IL-7 regardless of co-expression with the γc or not (**Fig. S1F-G**). Based on the above results and those previously reported by others^9,10^, we conclude that the disease mutation V253G of IL-7R can activate drives ligand-independent STAT5 activation comparable to IL-7 induced signaling by WT IL-7R in our expressed receptor system.

To investigate the structural effect of the V253G mutation by NMR, we reconstituted the IL-7R TMD with the point-mutation V253G (designated IL-7Rtm-V253G) in POPC-DH_6_PC bicelles (**Fig. S2**) that mimic a lipid bilayer^21–24^. To avoid biased dimerization due to disulfide bond formation, we substituted Cys261 by His, which is found at this position in mouse and rat IL-7R variants (**Fig. S3**). After the assignment of backbone amide and sidechain methyl chemical shifts (**Fig. S4),** we examined dimerization contacts by measuring NMR nuclear Overhauser effects (NOE). In this experiment, uniformly ^15^N-labeled and perdeuterated IL-7Rtm-V253G was mixed with ^13^C-labeled IL-7Rtm-V253G for exclusive detection of NOEs between amide protons of the (^15^N, ^2^H)-labeled chain with non-labile protons of the ^13^C-labeled chain^22^. The results show that the backbone amides of F250, V257, and A260 exhibited strong inter-chain NOEs (**Fig. 1A**). In comparison, the backbone amides of the WT IL-7Rtm (IL-7Rtm-WT) showing strong inter-chain NOEs are that of L255 and L259 (**Fig. 1B**). The change in inter-chain NOE pattern indicates that the V253G mutation resulted in about 170° rotation of the dominant dimerization interface from the L255 side to the G253 side (**Fig. 1C**). Subsequence full-scale analysis of intra- and inter- chain NOEs determined the high-resolution structure of the IL-7Rtm-V253G homodimer (**Fig. 1D and Table S1**).

The IL-7Rtm-V253G dimer structure is characterized by a right-handed helical packing mode with a packing angle of ∼30°. In the middle of the TMD, the S249-XXX-G253 sequence forms the most intimate inter-helical contact. The C-terminal ends of the two TMDs (residue K265) in the dimer are approximately 15 Å apart. The dimeric complex appears to be largely mediated by the small-XXX-small motif that is known to dimerize TM helix in many other single-pass membrane proteins^25–30^. The small-XXX-small motif, however, is not the only point of close helical packing as S249 also makes contact with the opposing helix in the dimer interface (**Fig. 1E**). Overall, the structure shows that replacing V253 with Gly has reshaped the face of the TMD involving residues 245, 249, 253, and 257 and turned it into a strongly dimerizing interface. According to our NOE data in **Fig. 1A**, this new face can dimerize stronger than the one involving residues 247, 251, 255, and 259.

**Fig. 1.**
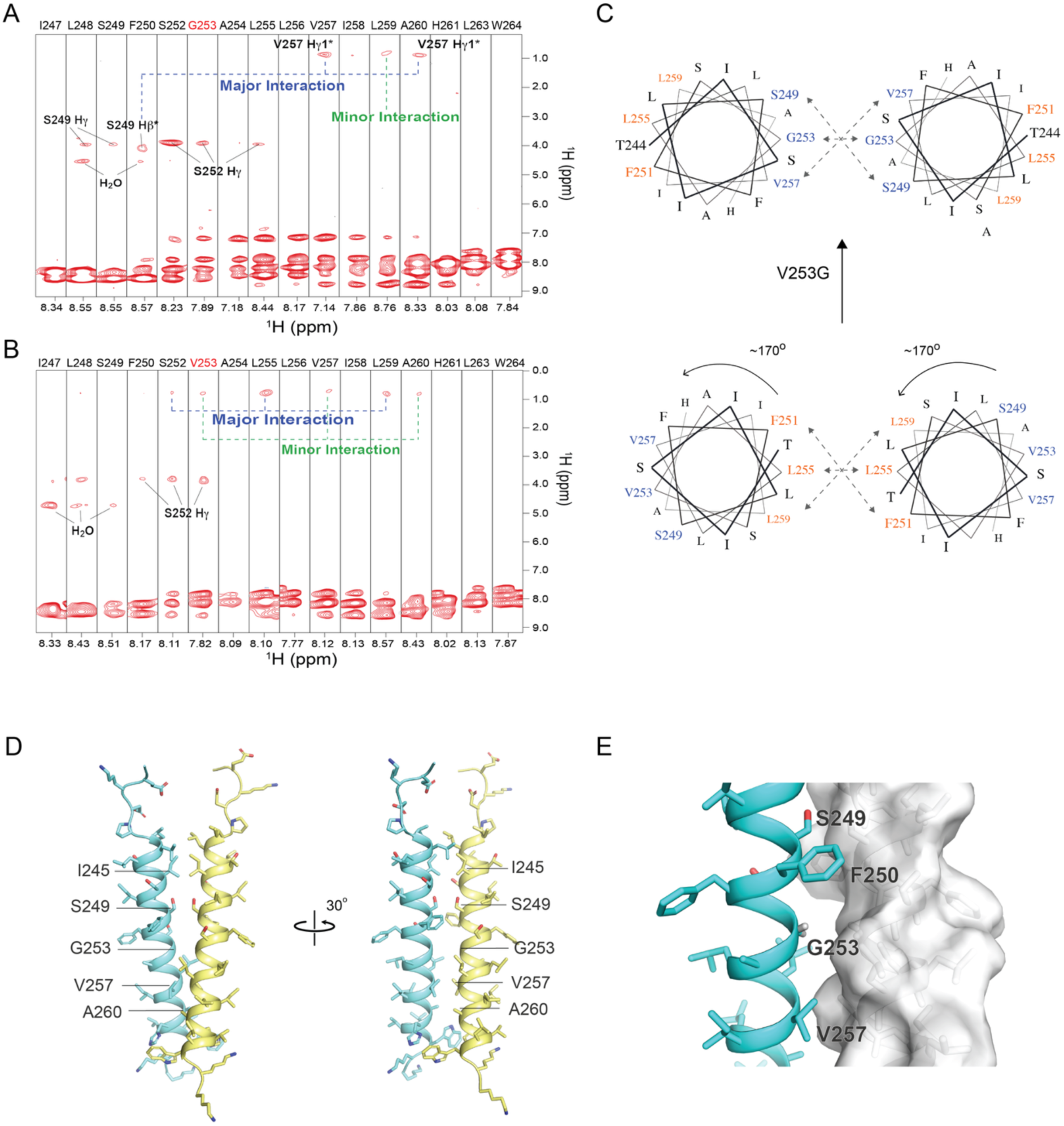
Major reorganization of the IL-7R TMD dimerization interface caused by the V253G gain-of-function mutation. A) Strips from 3D ^15^N-edited NOESY–TROSY spectra of (^15^N, ^2^H)-labeled IL-7Rtm-V253G randomly mixed with ^13^C-labeled IL-7Rtm-V253G showing interchain NOEs between the amide protons of (^15^N, ^2^H)-labeled chain and non-labile protons of the ^13^C-labeled chain. The spectra were recorded at 306 K at ^1^H frequency of 900 MHz. (B) Same type of NMR experiment as in (A) recorded with IL-7Rtm-WT at an 800 MHz spectrometer. (C) Helical wheel representations of the homodimeric interactions of IL-7Rtm-V253G (top) and IL-7Rtm-WT (bottom), showing that the V253G mutation resulted in ∼170° rotation of the helical packing interface. (D) Cartoon representation of the IL-7Rtm-V253G dimer structure. (E) Surface representation showing the dimer packing interface.

### Atomistic simulations identify altered stability and structural organization of TMD dimers

To investigate how the V253G mutation alters IL-7R TMD homodimerization, we generated models of the WT IL-7R TMD in addition to our NMR structure of the V253G mutant, which homodimerizes through the G253 interface (designated Mutant-G253; **Fig. 2A**). Wild-type structures were generated using two approaches: back-mutation of the V253G NMR structure while preserving the G253 interface (designated WT-V253; **Fig. 2B**), and Boltz-2 predicted structures guided by WT NMR constraints (**Fig. 1B**) corresponding to the L255 homodimerization interface (designated WT-L255; **Fig. 2C**). All dimers were embedded in POPC lipid bilayers with explicit solvent and ions for atomistic molecular dynamics (MD) simulations.

To compare the intrinsic stability of the dimer interfaces, we performed thermal stability simulations with the temperature gradually increased from 310 to 350 K over 200 ns (**Table S2**). Using the observed dissociation events from these simulations, we inferred dissociation temperatures as a measure of relative interface stability (**Fig. 2D**). Mutant-G253 was the most stable, with only one replicate dissociating across all simulations. WT-V253 was substantially less stable, with all replicates dissociating by ∼335 K. WT-L255 exhibited intermediate stability, with several replicates remaining associated at 350 K. Overall, these quantified stability profiles are consistent with experimental observations, with the V253G mutant being substantially more stable than WT variants.

To explore the conformational landscape and potential interconversion between interfaces, we performed extended MD simulations at 310 K for 1–2 μs per replicate, with 10 independent repeats per system (**Table S2**). Clustering analysis of intermolecular distances, reduced to two dimensions via principal component analysis (PCA), identified two compact, well-separated clusters along with dissociated states (**Fig. S5A**). The low within-cluster root-mean-square deviation (RMSD) compared to the higher between-cluster RMSD confirms that each cluster represents a distinct, well-defined conformational state (**Fig. S5B**). The first cluster, which we call the Mutant-like cluster, corresponds to the G253 mutant interface. The second cluster, which we call the WT-like cluster, corresponds to the L255 WT interface. Analysis of cluster populations showed very limited interconversion between interfaces (**Fig. 2E**). The Mutant-G253 simulations remained almost entirely as the G253 interface, with dissociation below 5%. The WT-V253 and WT-L255 simulations exhibited ∼10% and ∼13% dissociation, respectively.Both systems largely preserved their initial interface; only WT-V253 showed a minor transition toward the L255 interface (∼2%). The observation that the data resolve into just two clusters, each corresponding to an initial interface, with little interconversion between them, indicates that these two interfaces dominate the conformational landscape and are separated by an energetic barrier.

Contact analysis of the clustered data reveals the molecular basis for these distinct geometries (**Fig. 2F-H**). The Mutant-like cluster in Mutant-G253 and WT-V253 systems are stabilized by additional interactions at I245, S249, I248, L256, V257, and W264 (**Fig. 1A-B and Fig. 2F-G**). While the signature contact patterns of the Mutant-like interface are captured in both WT and mutant, the WT exhibits fewer contacting residues, likely because of the steric hindrance due to the V253 side chain (compare **Fig. 2F-G**). Notably, W264 sidechains on the intracellular (C-terminal) side of the TMD form aromatic stacking interactions that further stabilize this interface – a feature not captured in the NOE spectra (**Fig. 1B**) and absent in the L255 interface (**Fig. 2H**). In contrast, the WT-like cluster interface exhibits a distinct contact pattern involving I245, I248, S252, L255, L259, and V262, with prominent off-diagonal contacts, suggesting that this interface may be more malleable than the Mutant-like interface.

**Fig. 2.**
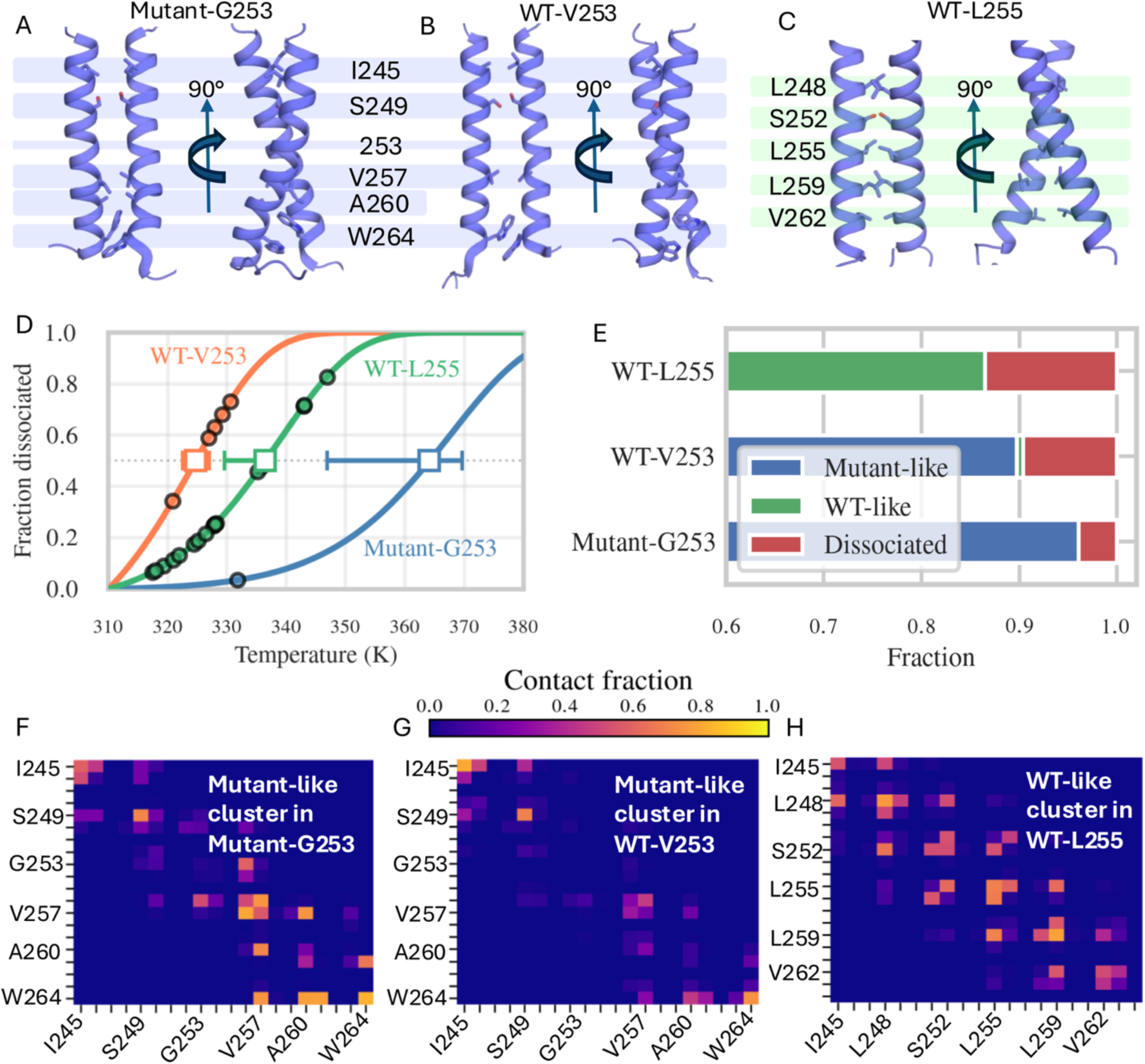
MD simulations uncover G253V-induced stabilization of IL-7R homodimers. **(A–C)** Cartoon representations of the Mutant-G253 and two WT systems (WT-V253 and WT-L255). The backbone is shown in cartoon and the main interfacial residues are highlighted in stick and labeled. **(D)** Thermostability of dimers from temperature-scan simulations. Survival curves were estimated assuming first-order kinetics with a temperature-dependent Arrhenius rate constant (see Methods). The dissociation temperatures (corresponding to 50% survival) are indicated by hollow squares, and error bars represent 95% confidence intervals. Data points (circles) show observed dissociation temperatures and are projected onto the survival curves. Even though no corresponding data points are shown, dimers that did not dissociate during the temperature scan (4/5 mutant NMR-based models, 4/21 WT modeled) still contribute to the survival curves and thereby inform the overall stability trends. **(E)** Fraction of replicates in each conformational cluster (interface type) or dissociated state, determined from clustering analysis of equilibrium MD simulations. **(F–H)** Contact frequency maps for Mutant-like and WT-like clusters captured in the Mutant-G253, WT-V253, and WT-L255 systems. The color bar shown in panel G applies to all maps.

Analysis of interhelical distance distributions in clustered data reveals distinct geometries for each interface **(Fig. S5C–E)**. The Mutant-like cluster, in both Mutant-G253 and WT-V253 systems, exhibits extracellular (N-terminal) and intracellular (C-terminal) distances centered around (8 Å, 10 Å), reflecting a relatively parallel helix arrangement. In contrast, the WT-like cluster shows a broader distribution, with the most probable conformation near (10 Å, 17 Å), consistent with a more prominent cross-like helix geometry. Because the intracellular separation determines the relative positioning of the JAKs, these results suggest that the mutation influences not only dimer stability but also its conformation in a manner that could support constitutive activation. Collectively, these MD simulations corroborate that the IL-7R TMD comprises two potential homodimerization interfaces on opposite sides, and that the V253G mutation induces a switch from the L255 to the G253 interface.

### TMD homodimerization at V253G is required for ligand-independent IL-7R signaling

We next implemented two independent biochemical experiments for validating the dimerization interface of the V253G mutant TMD. One experiment is based on a bacterial adenylate cyclase two-hybrid (BACTH) assay (**Fig. 3A**)^31^. Briefly, two complementary domains (T18 and T25) of adenylate cyclase (AC) were fused, separately, to the TMDs under investigation. The TMD fusion proteins were expressed in an AC deficient (CyaA) strain that yielded white bacteria colonies by default. Stable association of two TMDs reconstitutes AC activity, resulting in blue colonies. Another experiment, named TOXGREEN (**Fig. 3B**), is based on the expression of the super-folded Green Fluorescent Protein (sfGFP) reporter gene, which is driven by the dimerization of the bacterial transcription factor ToXR^32^. Specifically, ToXR and MBP (maltose binding protein) are fused to the N- and C- termini of the TMD, respectively, and the fusion protein (ToXR-TMD-MBP) is expressed in the inner membrane of *E. coli* with the MBP domain placed in the periplasm and the ToXR domain in the cytoplasm. Upon TMD dimerization, ToXR binds to the *ctx* promoter and activates the expression of sfGFP.

The results from both assays show that mutating S249 to Tyr on top of the base mutation V253G partially disrupted dimerization by 25-35% (**Fig. 3C-E**). Interestingly, replacing S249 with the smaller Ala, which was considered less disruptive, exhibited more disruptive power and reduced dimerization by 50-65% (**Fig. 3C-E**), suggesting that TMD dimerization is much more complicated than van der Waals (VDW) surface complementarity. Mutating V257 to Trp almost abolished TMD dimerization (**Fig. 3C-E**), consistent with the dimer structure. These mutagenesis results indicate that the dimer interface of IL-7Rtm-V253G involving S249, G253, and V257 revealed by the NMR structure mediates the homotypic association of IL-7Rtm-V253G in cell membrane.

To determine whether the V253G dimer interface is relevant to ligand-independent IL-7R signaling, we evaluated the effect of the above dimer-breaking mutations on signaling. Full-length WT IL-7R, IL-7R-V253G, IL-7R-V253G/S249Y, IL-7R-V253G/S249A and IL-7R-V253G/V257W with or without γc were expressed in BaF3 cells. Flow cytometry analysis showed that the membrane expression levels of these IL-7R variants were nearly the same (**Fig. S6**). Immunoblotting results show that the V253G mutation generated very significant ligand-independent p-STAT5 and p-JAK1 signals (**Fig. 3F-H**). The S249Y mutation on top of V253G caused about 50% and 30% reduction in p-STAT5 and p-JAK1, respectively. The S249A mutation reduced p-STAT5 and p-JAK1 signals more than S249Y (**Fig. 3F-H**), consistent with the TMD dimerization assays (**Fig. 3C-E**). The V257W mutation on top of V253G also reduced the levels of p-STAT5 and p-JAK1 (**Fig. 3F-H**). Overall, the results show clear correlation between disruption of V253G TMD dimerization and reduced levels of p-STAT5 and p-JAK1 signals, indicating that IL-7R TMD dimerization at the G253 interface is responsible for the observed ligand-independent JAK1 and STAT5 phosphorylation.

**Fig. 3.**
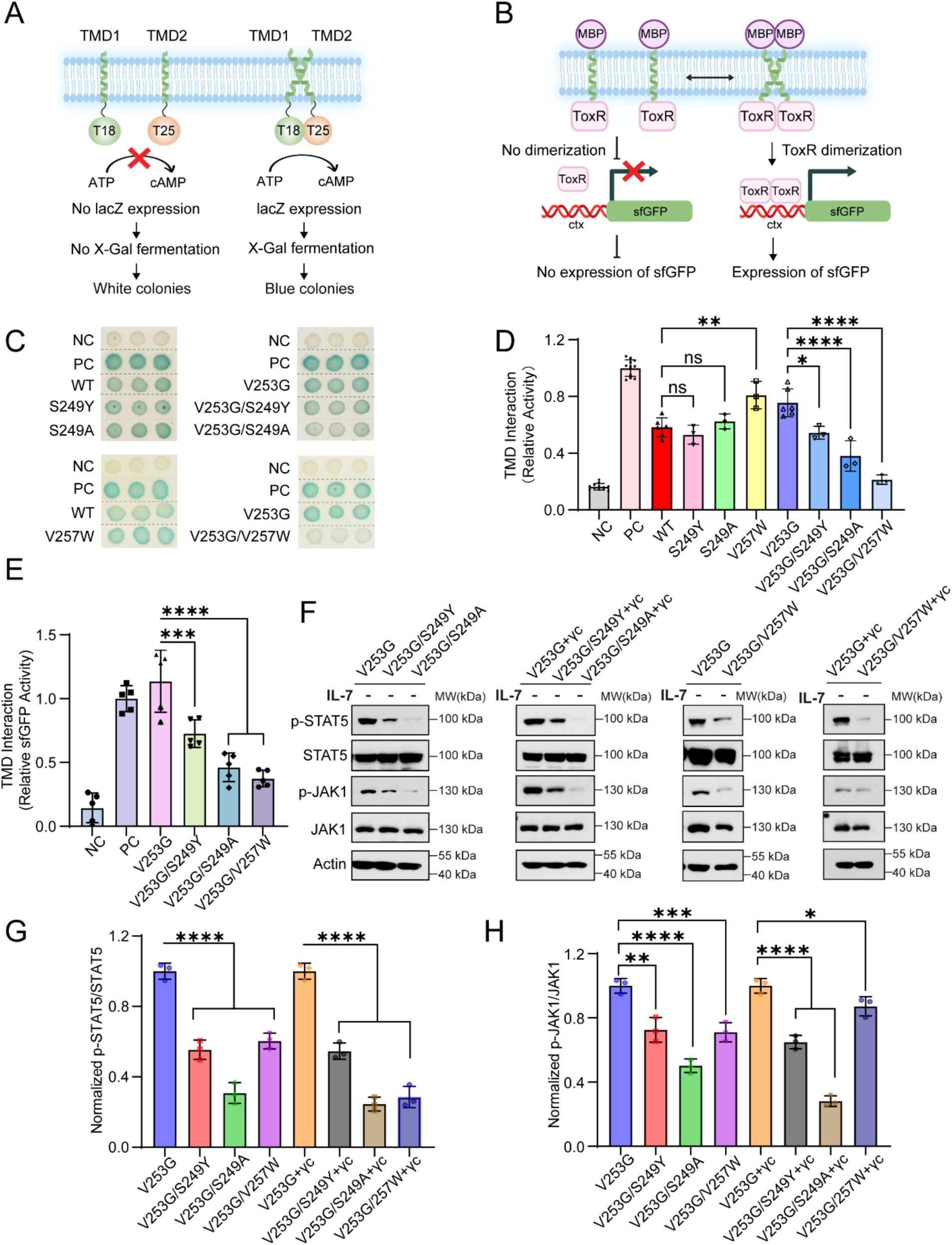
Requirement of TMD dimerization at the V253G interface by ligand-independent IL-7R signaling. (**A** and **B**) Schematic illustrations of BACTH and TOXGREEN analyses, respectively, used for examining homotypic interactions of IL-7R TMD variants. For the BACTH assay, IbaG/IbaG homodimerization was used as the positive control. For the TOXGREEN assay, glycophorin A (GpA), a classical TM dimer, was used as a positive control, while GpA-G83I, a monomeric mutant, served as the negative control. (C) BACTH analysis of TMD interactions for IL-7R TMD variants. Blue colonies indicate interaction between the TMDs in the bacterial inner membrane, while white colonies indicate no such interaction. (D) Quantification of BACTH colony colors normalized relative to the IbaG/IbaG positive control. The data are shown as means ± SEMs calculated from technical replicates (typically three) from three independent experiments. Statistical significance by One-way ANOVA. ns, not significant; \**p* ≤ 0.05; ** *p* ≤ 0.01; *** *p* ≤ 0.001; **** *p* ≤ 0.0001. (E) TOXGREEN analysis of TMD interactions for IL-7R TMD variants. Quantification of sfGFP activity is normalized relative to the GpA positive control. The data are shown as means ± SEMs calculated from technical replicates (typically five to six) from three independent experiments. Statistical significance by One-way ANOVA. ns, not significant; \**p* ≤ 0.05; ** *p* ≤ 0.01; *** *p* ≤ 0.001; **** *p* ≤ 0.0001. (F) Phosphorylation of STAT5 (p-STAT5) and JAK1 (p-JAK1) in BaF3 cells expressing WT or mutant IL-7R or co-expressing with γc. The p-STAT5 and p-JAK1 signals were detected by immunoblotting and compared with total STAT5 and JAK1, respectively. (**G-H**) Quantification of p-STAT5 and p-JAK1 signals in (F) as p-STAT5/STAT5 and p-JAK1/JAK1 intensity ratios, normalized to the V253G or V253G/γc. The data are shown as means ± SEMs calculated from three independent experiments. Statistical significance by One-way ANOVA. ns, not significant; \**p* ≤ 0.05; ** *p* ≤ 0.01; *** *p* ≤ 0.001; **** *p* ≤ 0.0001.

The above results have been independently validated by the STAT5 luciferase reporter assay. In this assay, a BaF3 cell line stably expressing a STAT5-responsive luciferase reporter was used to monitor the STAT5 signaling. Consistent with the immunoblotting data, all three double-point mutants showed significantly reduced luciferase activity (**Fig. S7**).

### The V253G mutation causes ligand-independent dimerization of IL-7R in live cells

To investigate the relevance of V253G-induced IL-7R TMD dimerization in the physiological context, we quantified receptor diffusion and interaction in the plasma membrane of live cells by single-molecule imaging techniques. IL-7R fused to an N-terminal ALFA tag (ALFA-IL-7R) was constructed, enabling highly selective cell surface labeling using anti-ALFA tag nanobodies (ALFAnb) conjugated with photostable fluorophores (**Fig. 4A**). Activity analysis by phospho-flow cytometry confirmed substantially increased p-STAT5 phosphorylation activity of ALFA-IL-7R V253G compared to WT (**Fig. S8A**). For single-molecule imaging, cells were complemented with JAK1 lacking the tyrosine kinase (TK) domain and fused to a C-terminal mEGFP (JAK1-ΔTK-mEGFP), to ensure receptor integrity while avoiding bias caused by downstream signal activation. HeLa cells transiently expressing ALFA-IL-7R and JAK1-ΔTK-mEGFP were labeled in *situ* by adding a mixture of ALFAnbs conjugated with donor and acceptor dye, respectively.

**Fig. 4.**
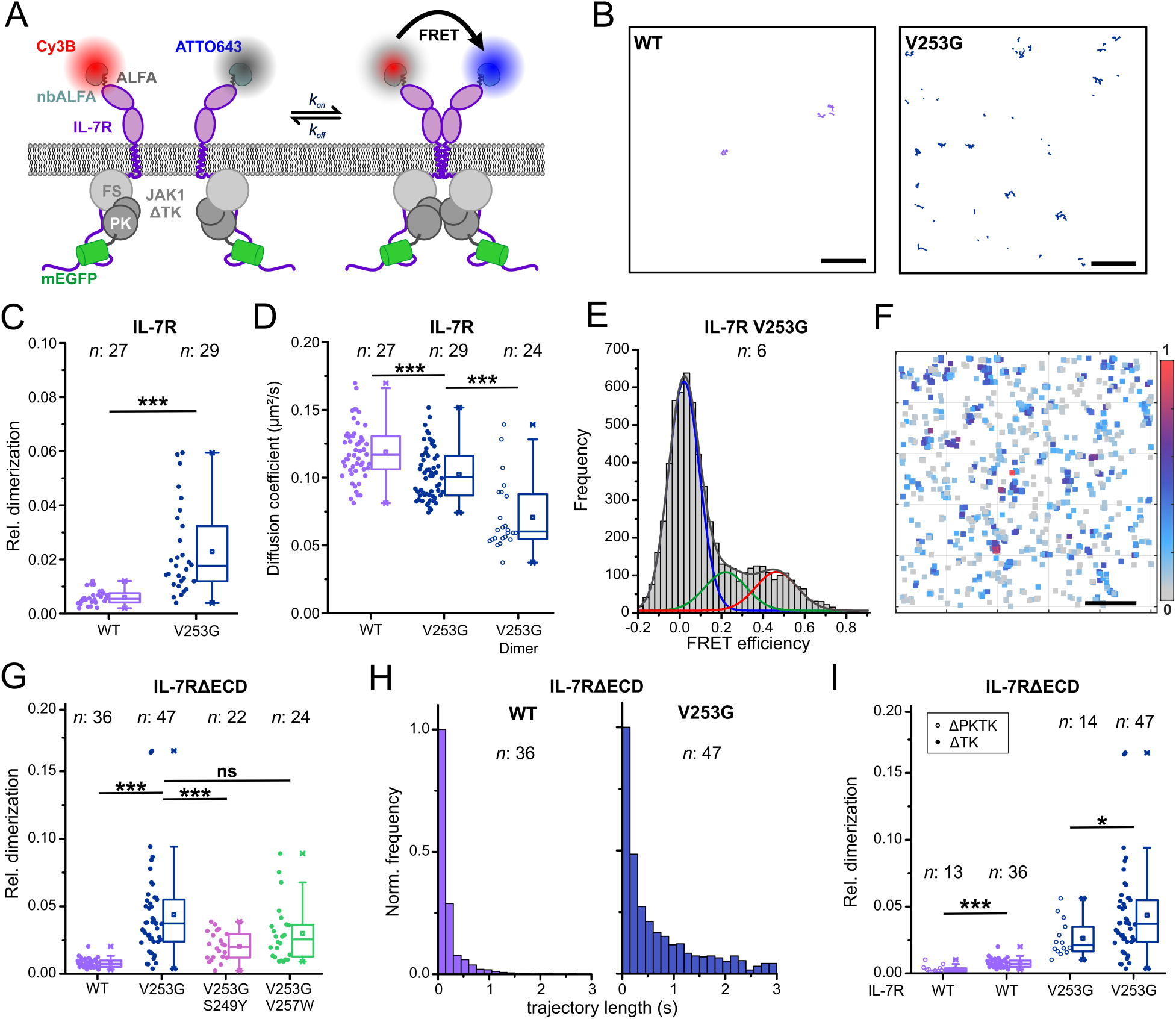
Ligand-independent dimerization of IL-7R in the plasma membrane of live cells. (A) In *situ* detection of IL-7R V253G dimerization by single-molecule FRET (smFRET) in live cells. IL-7R fused to an N-terminal ALFA-tag was expressed in HeLa cells and labeled in situ with anti-ALFA nanobodies conjugated with donor fluorophore Cy3B or acceptor fluorophore ATTO 643. Dimerization was detected by ALEX-FRET TIRF imaging. (B) Representative smFRET trajectories observed for IL-7R WT and V253G, respectively. Scale bar: 5 µm. (C) Dimerization of IL-7R WT and V253G quantified by smFRET. Data from one representative experiment, with each data point corresponding to the result from one cell. (D) Diffusion constants obtained from IL-7R WT and V253G donor and acceptor channel trajectories (solid circle), and V253G-induced co-trajectories (FRET-channel; open circle). Data from one representative experiment, with each data point corresponding to the result from one of *n* cells. (E) smFRET efficiency histogram of all co-localized IL-7R V253G pooled from *n* = 6 cells, and fit by multiple Gaussian functions. The peak with a FRET efficiency of ∼0 (blue fit curve) corresponds to randomly co-localized donor and acceptor molecules. (F) Localization map of co-localized donor and acceptor signals from 150 consecutive frames, color-coded according to the corresponding FRET efficiency. Scale bar: 5 µm. (G) Comparison of dimerization levels for different IL-7RΔECD variants obtained from co-tracking and smFRET detection. Data from three independent experiments, with each data point corresponding to the result from one of *n* cells. (H) smFRET trajectory length histograms for IL-7RΔECD WT and V253G dimers from one representative experiment pooled from *n* cells (>1000 trajectories for each condition). (I) Dimerization of IL-7RΔECD V253G in the presence of different JAK1 variants. Data from one representative experiment, with each data point corresponding to the result from one of *n* cells. Statistical significance in **C**, **D**, **G** and **I** by Student’s t test. ns, not significant; \**p* ≤ 0.05; ** *p* ≤ 0.01; *** *p* ≤ 0.001; **** *p* ≤ 0.0001.

Interrogation by total internal reflection fluorescence (TIRF) microscopy with alternating laser excitation (ALEX)^32^ revealed individual IL-7R at total densities < 2/µm², mostly randomly diffusing in the plasma membrane (**Fig. S8B and Movie S1**). For WT IL-7R, very few, short-lived single-molecule Förster resonance energy transfer (smFRET) events were observed, confirming very low homodimerization propensity (**Fig. 4B and Movie S1)**. IL-7R V253G, however, showed significantly higher level of dimerization, with longer smFRET trajectories (**Fig. 4B and Movie S1**). Quantitative co-tracking and smFRET trajectory length analyses confirmed enhanced dimerization levels of V253G compared to WT IL-7R and increased dimer stability (**Fig. 4C and Fig. S8C**). Diffusion constants obtained by mean squared displacement (MSD) analysis from single-molecule trajectories confirmed ∼40% reduced mobility for V253G-induced IL-7R dimers as compared to the average diffusion of most monomeric IL-7R WT (**Fig. 4D**), in line with previous observations for other class I/II cytokine receptor dimers^20,33,34^. Quantitative smFRET efficiency analysis for IL-7R V253G dimers yielded a distribution with peaks at 22% and 46% (**Fig. 4E**). This may either indicate a broad distribution of orientations due to high flexibility of the ectodomains or a mixture of two conformers. Localization maps of co-localized IL-7R V253G suggest that the latter may be the case, as trajectories with distinct smFRET efficiencies can be discerned (**Fig. 4F**). However, decisive analysis of potential conformational dynamics is obstructed by the relatively few datapoints within individual trajectories.

For detecting the weak, transient dimerization more robustly, we further increased the FRET efficiency by using IL-7R with its ectodomain replaced by a non-fluorescent monomeric enhanced green fluorescent protein (mXFP-IL-7RΔECD), which was labeled via anti-GFP nanobodies (**Fig. S8D**)^35^. Activity assays by phospho-flow cytometry showed similar responsiveness to the V253G mutation for the mXFP-IL-7RΔECD construct as compared to ALFA-IL-7R (**Fig. S8A**). In HeLa cells co-expressing mXFP-tagged IL7RΔECD or IL-7R-V253GΔECD together with JAK1-ΔTK-mEGFP, V253G exhibited substantially higher dimerization than WT (**Fig. 4G, Fig. S8E and Movie S2).** The much higher FRET efficiency of ∼0.8 enabled tracking of smFRET signals with high fidelity (**Fig. S8F and Movie S2**). Thus, dimer stabilities could be more robustly determined by smFRET trajectory length analysis, confirming substantially increased dimer stability for V253G as compared to WT (**Fig. S8H-I**). Quantitative co-tracking and smFRET trajectory length analyses confirmed enhanced dimerization levels and dimer stability for V253G as compared to the WT (**Fig. 4C-D and Fig. S8B**). The mutation S249Y significantly weakened V253G-induced dimerization of IL-7RΔECD and reduced the lifetime of dimers, while V257W showed a weaker but non-significant reduction (**Fig. 4G and Fig. S8G-I**), in line with their reduced activity in functional studies (**Fig. 3F-H**).

**Fig. 5.**
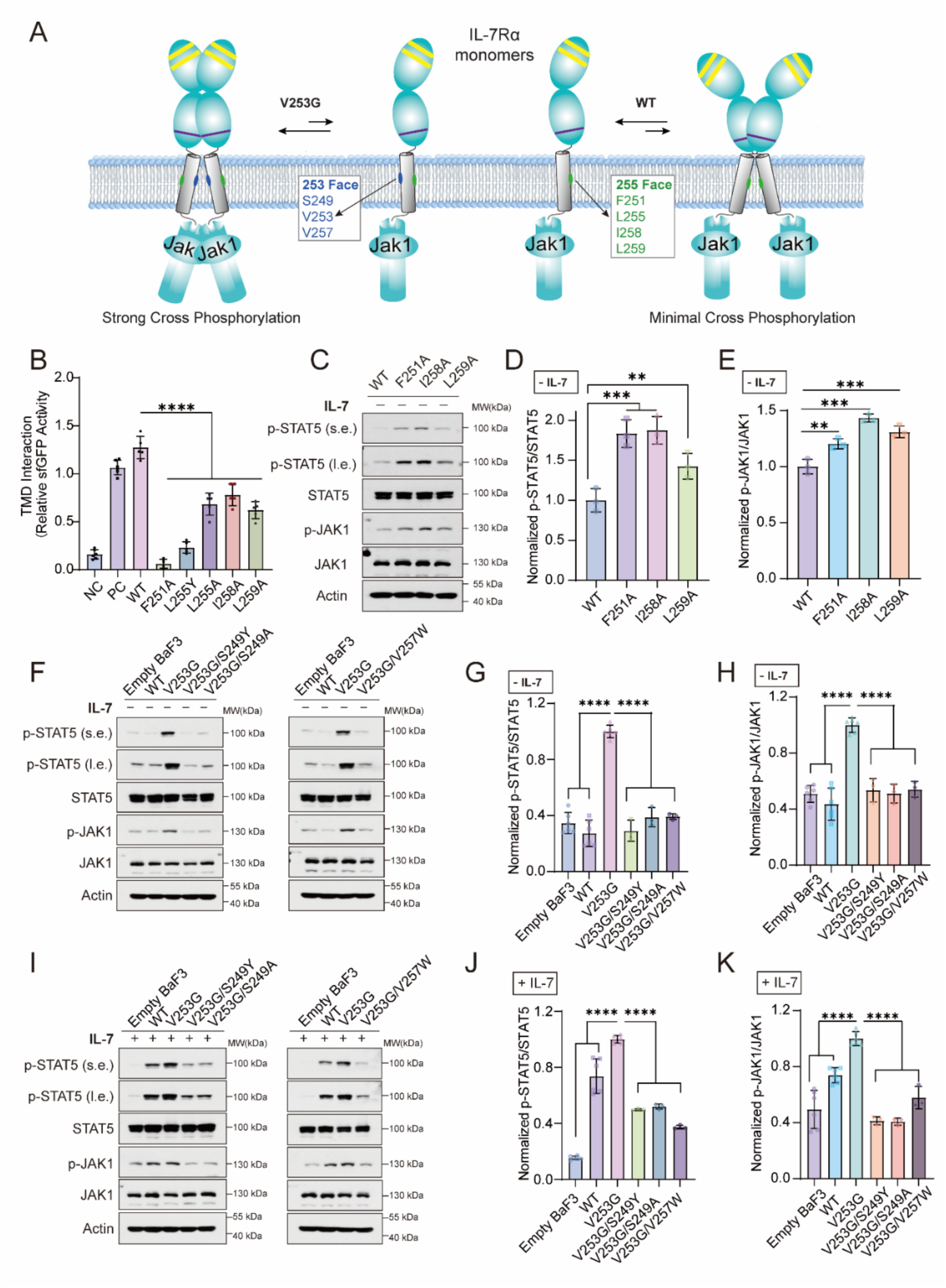
Activating homodimer of V253G IL-7R vs. inactivating homodimer of WT IL-7R. (A) Schematic illustration of activating homodimer of V253G IL-7R and inactivating homodimer of WT IL-7R. In the V253G IL-7R, residues S249, V253, and I257 form the homodimerization interface, which promotes a JAK1–JAK1 pairing configuration that leads to strong cross phosphorylation. In contrast, the WT IL-7R forms a homodimerization interface composed of F251, L255, I258, and L259, which is incompatible with JAK1-JAK1 cross phosphorylation. (B) TOXGREEN analysis of the impact of single mutations at the L255 face on IL-7R-TMD dimerization. Quantification of sfGFP activity is normalized relative to the GpA positive control. The data are shown as means ± SEMs calculated from technical replicates (typically five to six) from three independent experiments. (C) Effect of disrupting the L255 dimerizing face on ligand-independent p-STAT5 and p-JAK1 signals in BaF3 cells. After 5 h of cytokine deprivation, cells expressing WT or mutant IL-7R (F251A, F258A, F259A) were treated with PBS. The p-STAT5 and p-JAK1 signals were detected by immunoblotting and compared with total STAT5 and JAK1, respectively. (**D-E**) Quantification of p-STAT5 and p-JAK1 signals in (C) as p-STAT5/STAT5 and p-JAK1/JAK1 intensity ratios, normalized to the WT IL-7R. (**F**) Effect of disrupting the G253 dimerizing face on ligand-independent p-STAT5 and p-JAK1 signals in BaF3 cells. After 5 h of cytokine deprivation, cells expressing WT or mutant IL-7R (V253G, V253G/S249Y, V253G/S249A, V257W) were treated with PBS. (**G-H**) Quantification of p-STAT5 and p-JAK1 signals in (F) as p-STAT5/STAT5 and p-JAK1/JAK1 intensity ratios, normalized to the V253G IL-7R. (**I**) Effect of disrupting the G253 dimerizing face on ligand-dependent p-STAT5 and p-JAK1 signals in BaF3 cells. After 5h of cytokine deprivation, cells expressing WT or mutant IL-7R (V253G, V253G/S249Y, V253G/S249A, V257W) were treated with 50 ng/mL IL-7 (**J-K**) Quantification of p-STAT5 and p-JAK1 signals in (I) as p-STAT5/STAT5 and p-JAK1/JAK1 intensity ratios, normalized to the V253G IL-7R.

To explore potential cooperation between TMD and JAK1 homotypic interactions, we compared dimerization of IL-7RΔECD WT and V253G in the presence of JAK1-ΔTK as well as JAK1 lacking both TK and pseudokinase (PK) domains (JAK1-ΔPKTK). Previous structural and functional studies have shown that productive interaction between the PK domains is relevant for releasing the TK domain and initiate downstream signaling^19,20,36^. Strikingly, we found significantly reduced dimerization of IL-7RΔECD V253G upon co-expression with JAK1-ΔPKTK (**Fig. 4I**), highlighting that the V253G mutation promotes a TMD dimer geometry compatible with self-association of the JAK1 PK domain. Interestingly, IL-7RΔECD WT showed a similar effect although dimerization of the WT was very low to begin with.

Taken together, our smFRET results demonstrate the new dimerization interface induced by the V253G mutation in live cells and its synergy with JAK1 dimerization between the PK domains, which is important for signal activation.

### Distinct activating and inactivating homodimerization interfaces of the IL-7R TMD

The IL-7Rtm-WT at high concentration shows specific intermolecular NOEs on the L255 face of the TMD (**Fig. 1**), consistent with the MD and biochemical analyses that the WT TMD by itself can homodimerize (**Fig. 2 and 3**). It is unclear, however, whether this mode of homotypic interaction has any functional consequences. We thus compared the effects of mutating the G253 interface of the V253G mutant and the L255 interface of the WT on ligand-independent receptor signaling. Based on the observation that the V253G mutant but not the WT shows strong signaling in the absence of IL-7 (**Fig. S1F-G**), we hypothesized that TMD homodimerization at the G253 face is structurally compatible with JAK1-JAK1 cross phosphorylation whereas TMD homodimerization of the WT at the L255 face is not (**Fig. 5A**). We first examined the effect of single mutations at the L255 face (F251A, L255Y, L255A, I258A, L259A) and found that disrupting the L255 interface indeed reduced dimerization level of IL-7R-TMD (**Fig. 5B**). We then tested the effect of these mutations on STAT5 and JAK1 phosphorylation in BaF3 cells. In the absence of IL-7, single mutations expected to disrupt the packing at the L255 interface – F251A, I258A, and L259A – all exhibited substantially higher levels of p-STAT5 and p-JAK1 than that of the WT (**Fig. 5C-E**). These unexpected results suggest that TMD dimerization at the L255 face is incompatible with JAK1-JAK1 cross phosphorylation, as depicted in **Fig. 5A**.

For a direct comparison with the V253G mutant, we also tested single mutations S249Y, S249A, and V257W on top of the V253G mutation that alter the G253 dimerization interface. These mutations all showed dramatic reduction in p-STAT5 and p-JAK1 levels in the absence of IL-7 (**Fig. 5F-H**). The immunoblotting results overall agree with the smFRET observation that S249Y significantly decreased the mutant dimerization levels on the cell surface (**Fig. 4G**).

In all quantification graphs, data are shown as means ± SEMs calculated from three independent experiments. Statistical significance by One-way ANOVA. ns, not significant; \**p* ≤ 0.05; \*\**p* ≤ 0.01; \*\*\**p* ≤ 0.001; \*\*\*\**p* ≤ 0.0001. PBS, phosphate-buffered saline. (s.e.) = short exposure. (l.e.) = long exposure.

Addition of IL-7 could increase the level of WT receptor signaling, due to ligand-induced pairing with endogenously expressed γc, but not that of the single mutant V253G and double mutants V253G/S249Y, V253G/S249A, and V253G/V257W (**Fig. 5I-K**). Thus, these mutations, despite of being on the opposite side of the L255 interface recognized by the γc TMD^18^, could allosterically weaken IL-7R TMD interaction with the γc TMD.

**Fig. 6.**
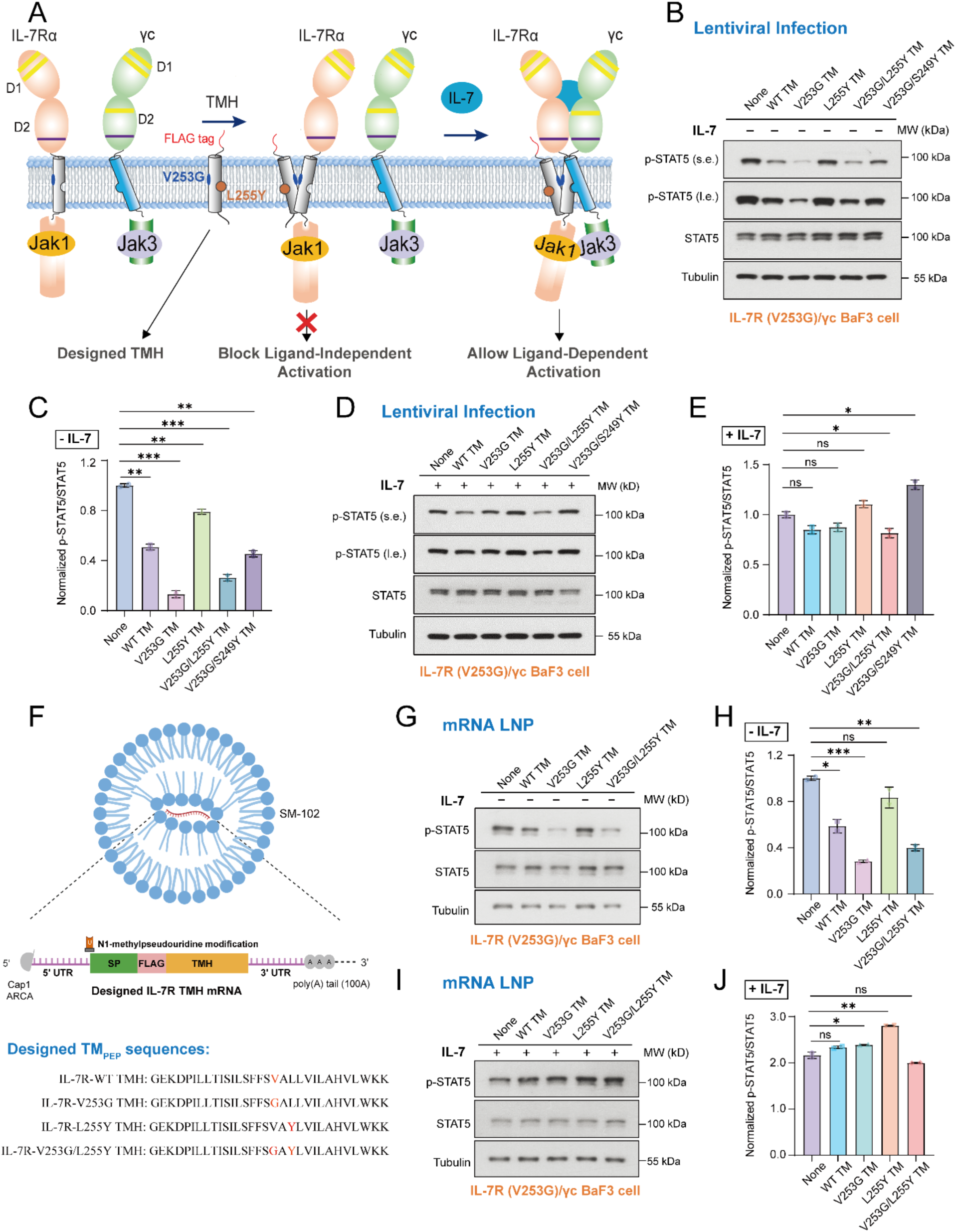
Selective interference with the oncogenic IL-7R signaling via rational helices design. (A) Schematic illustration of the TMD design concept for competitive blocking of IL-7R V253G homodimerization and ligand-independent receptor activation but not effecting IL-7 induced signaling. (B) Immunoblot analysis of STAT5 phosphorylation (p-STAT5) in stable BaF3 cells expressing the IL-7R V253G mutant and γc, following lentiviral transduction with WT TM_PEP_ or mutants (V253G, L255Y, V253G/L255Y, V253/S249Y). After 12 h of cytokine deprivation, cells were treated with PBS and collected for immunoblotting. (C) Quantification of p-STAT5 signal in (B) as p-STAT5/STAT5 intensity ratios, normalized to the no-delivery control (None). (D) Immunoblot analysis of p-STAT5 in stable BaF3 cells expressing the IL-7R V253G mutant and γc, following lentiviral transduction with WT TM_PEP_ or mutants as in (B). After 12 h of cytokine deprivation, cells were treated with 50 ng/mL IL-7 for 2 h and collected for immunoblotting. (E) Quantification of p-STAT5 signal in (D) as p-STAT5/STAT5 intensity ratios, normalized to the no-delivery control (None). (F) Schematic illustration of the mRNA–LNP delivery system. The mRNA encoding designed TM_PEP_ was delivered into BaF3 cells using lipid nanoparticles (LNPs) comprising ionizable lipids (SM-102) and auxiliary components (*Top*). The mRNAs were modified with N1-methylpseudouridine (m1Ψ) and contained a 5′ Cap1/ARCA and 3′ poly(A) tail (∼100A) (*Middle*). The designed TM_PEP_ sequences were listed with mutated residues in red (*Bottom*). (G) Immunoblot analysis of p-STAT5 in stable BaF3 cells expressing the IL-7R V253G mutant and γc, following mRNA**–**LNP delivery of WT TM_PEP_ or mutants (V253G, L255Y, V253G/L255Y, V253/S249Y). After 12 h of cytokine deprivation, cells were treated with PBS and collected for immunoblotting. (H) Quantification of p-STAT5 signal in (G) as p-STAT5/STAT5 intensity ratios, normalized to the no-delivery control (None). (I) Immunoblot analysis of p-STAT5 in stable BaF3 cells expressing the IL-7R V253G mutant and γc, following mRNA**–**LNP delivery of WT TM_PEP_ or mutants as in (G). After 12 h of cytokine deprivation, cells were treated with 50 ng/mL IL-7 for 2 h and collected for immunoblotting. (J) Quantification of p-STAT5 signal in (I) as p-STAT5/STAT5 intensity ratios, normalized to the no-delivery control (None). Unpaired student’s t test was used and data were shown as mean ± SEM calculated from three independent experiments. Represents statistically significant, n.s. (not significant); \**p* ≤ 0.05, \*\**p* ≤ 0.01, \*\*\**p* ≤ 0.001, \*\*\*\**p* ≤ 0.0001. PBS, phosphate-buffered saline. s.e.=short exposure. l.e.= long exposure.

Collectively, the above results suggest that the dimerization mode of the IL-7R TMD can have profound influence on receptor activation. TMD dimerization at the G253 face can directly activate intracellular JAK1/STAT5 signaling without ligand binding. In contrast, TMD dimerization at the L255 face appears to prevent random JAK1-JAK1 cross activation and such autoinhibitory mechanism may be important in scenarios of high receptor expression level and high frequency of ligand-independent IL-7R dimerization.

### Designed TM helices selectively block ligand-independent but not ligand-dependent signaling

The strong NOEs observed at the homodimer interface of the V253G mutant (**Fig. 1A**) suggest that this non-native, disease-causing interaction is specific. More importantly, the structural analysis indicated that the non-native dimerization does not compete with interaction with the γ-chain TMD, thereby affording the opportunity to design TM peptide (TM_PEP_) to block the disease-causing homodimerization without affecting the native heterodimerization. We then explored the use of a TM_PEP_ based on the IL-7R TMD sequence, which can compete with TMD homodimerization of the V253G mutant but lacks the receptor ICD necessary for signaling, thus reversing the aberrant signaling caused by the TM mutation. Specifically, the TM_PEP_ contained the V253G mutation for blocking IL-7R mutant homodimerization, or ligand-independent signaling, while simultaneously carrying an L255Y mutation for disrupting the pocket that interacts with the γc TMD, ensuring that it does not interfere with ligand-dependent activation (**Fig. 6A**). A signal sequence was added to the N-terminus of the TM_PEP_ to direct it to the plasma membrane upon synthesis. Moreover, a FLAG-tag after the signal sequence was included for probing proper membrane insertion of the TM_PEP_ with anti-FLAG antibody.

To test the above hypothesis, we used two different approaches to deliver designed TM_PEP_ to BaF3 cells: one is lentiviral infection with DNA encoding the TM_PEP_; in another method, the mRNA coding for the TM_PEP_ was delivered using lipid nano particle (LNP). For lentiviral infection, the WT TM_PEP_ or mutants (V253G, L255Y, V253G/L255Y, V253/S249Y) were cloned into lentiviral vector with a signal sequence and a FLAG-tag on the N-terminus. Following lentiviral infection of BaF3 cells expressing the IL-7R V253G mutant and the WT γc, flow cytometry selection using anti-FLAG antibody was performed to generate cell lines stably expressing the TM_PEP_ variants. The cell surface expression levels of the IL-7R V253G mutant and the WT γc were not significantly affected by the transfection of various designed TM_PEP_ (**Fig. S9**).

In the absence of IL-7 (**Fig. 6B-C**), the V253G mutant again exhibited very robust STAT5 phosphorylation. Upon co-expressing the TM_PEP_ with either single mutation V253G or double mutations V253G/L255Y, which are expected to block homodimerization of the IL-7R V253G mutant, the ligand-independent p-STAT5 signal was drastically reduced by ∼70%. Introducing the S249Y on top of the V253G in the TM_PEP_ resulted in weaker inhibition of ligand-independent signaling, consistent with the observation that S249Y can partially disrupt V253G-mediated TMD dimerization. The TM_PEP_ baring the single mutation L255Y did not show significant inhibition because L255 is not directly involved in dimerization at G253. Interestingly, co-expressing the WT TM_PEP_ also resulted in partial but significant inhibition of tonic signaling (**Fig. 6B-C**), which is likely due to the minor homodimerization interface on the V253 side of the WT TMD (**Fig. 1B**). In the presence of IL-7, however, co-expression of none of the TM_PEP_ variants showed significant inhibitory effect on STAT5 phosphorylation (**Fig. 6D-E**), likely because the TM_PEP_ cannot compete with the stronger ligand-induced TM heterodimerization of the full-length IL-7R and γc

In addition to lentiviral infection, we tested the delivery of TM_PEP_ more directly using mRNA encapsulated in LNP, which has been a technology successfully used for vaccine delivery.^37^ The mRNA encoding WT TM_PEP_ or mutants (V253G, L255Y, V253G/L255Y, V253/S249Y) were chemically modified with N1-methylpseudouridine (m1Ψ) and contained a 5′ Cap1/ARCA and 3′ poly(A) tail (∼100A)(**Fig. 6F**). These mRNAs were encapsulated in LNPs composed of SM-102, along with auxiliary components essential for nanoparticle stability and efficient delivery. The aforementioned engineered BaF3 stable cells expressing the IL-7R V253G mutant and the γc were starved without cytokines overnight. The starved cells were treated with different mRNA-LNPs and harvested for immunoblotting to examine the effects on downstream JAK/STAT signaling. The results from administration of mRNAs encoding the designed TM_PEP_ are highly consistent with that of the lentiviral infection method, especially in the absence of IL-7 (**Fig. 6G-J**).

## DISCUSSION

Naturally occurring GOF mutations within TM regions of single-pass TM receptors have been observed in many types of cytokine and growth factor receptors^1–4,6,10^. The mechanism of action for these TM mutations has been elusive due to insufficient structural information for the TM and membrane-proximal regions of these receptors as well as the interplay between the TMD and the intracellular signaling domain that allow a TM mutation to activate downstream signaling completely independent of extracellular ligand binding. In this study, we find that the GOF disease mutation V253G residing in the TMD of IL-7R acts by structural rewiring of IL-7R homodimerization leading to intracellular JAK1-JAK1 activation independent of the ligand.

Specifically, the V253G mutation resulted in a major shift of the homotypic TMD dimerization from the L255 interface of the WT (F251, L255, L259) to the G253 interface of the mutant (S249, G253, V257) (**Fig. 1 and 2**). Interestingly, cysteine substitutions found in the IL-7R TMD from ALL patients are predominantly at residue positions 242 and 249^9,11^, which are both predicted to crosslink the G253 interface, consistent with the activating TMD dimerization of the V253G mutant. The G253 and L255 homodimerizing interfaces are opposite of each other on the TMD. They also have opposite signaling consequences, as disrupting the G253 interface resulted in drastic reduction of ligand-independent STAT5 and JAK1 phosphorylation whereas breaking the L255 interface increased ligand-independent signaling (**Fig. 5C-H**). These results suggest that mere TMD dimerization is insufficient for activating intracellular signaling. The TMD needs to dimerize in a specific manner in order to position the JAK1s in an arrangement suitable for initiating cross phosphorylation. Although the structural link between the TMD and JAK1 is unclear, our smFRET results (**Fig. 4I**) strongly suggest that dimerization through the G253 interface enables productive conformational coupling between the TMD and the PK domain of JAK1 and thus signal activation.

The incompatibility between the L255 interface homodimerization and JAK1 activation could represent a previously unrecognized function of TMD dimerization in receptor control and autoinhibition. Homodimerization of the WT IL-7R TMD via the L255 interface has been observed in the NMR and BACTH analyses, although both showed that it is weaker than the G253 interface dimerization of the mutant TMD (**Fig. 1A-B and 3C-D**). The NMR and BACTH experiments, however, were performed at artificially high protein concentration. In the smFRET experiment where the receptor concentration was kept low for single-molecule tracking, the WT showed insignificant dimerization but the V253G mutant did (**Fig. 4B-C**). Consistently, MD simulations confirm that the G253 dimer interface is severely destabilized upon re-introducing V253 (**Fig. 2D**). Our previous NMR study also showed that when mixed with the TMD of γc, the L255 interface homodimerization cannot compete with its heterodimerization with the γc TMD^18^. The simulations demonstrate that the V253G mutation stabilizes a parallel helix geometry, characterized by similar extracellular (N-terminal) and intracellular (C-terminal) spacings reinforced by W264-mediated aromatic stacking. In contrast, the WT dimer with the L255 interface adopts a crossed-helix arrangement with larger separation at both the extracellular and intracellular sides and lacks W264 stabilization. These structural differences have direct implications for JAK1 positioning and receptor activation, as the reinforced intracellular spacing in the V253G mutant likely enforces an optimal distance and orientation for constitutive JAK1-JAK1 interaction. It is tempting to speculate that the other co-receptor of IL-7R, CRLF2, which also has a Trp residue upstream of the capping Lys (W252), may be recruited via the V253 interface. Interestingly, CRLF2 signals via JAK2, which is more similar to JAK1 than to JAK3; its juxtamembrane segment, which is important for defining functional JAK coupling, is much more similar to IL-7R than to γc. The above clues collectively suggest that the homotypic interaction of the IL-7R TMD at the L255 face is conformationally specific but intrinsically weak. As such, this dimeric interaction may serve as a safety for preventing ligand-independent JAK1 phosphorylation at high receptor concentrations, which can be easily released in the presence of ligand and γc. This safety mechanism, however, cannot prevent dimerization of the G253 interface of the V253G mutant that causes constitutive JAK1 activation. A similar switch of transmembrane dimer interfaces has been proposed to be involved in EGFR activation^28,38,39^, yet experimental validation in the cellular context has remained controversial^40^.

Directly targeting receptor TMD with antibody and small molecule is technically challenging due to additional consideration of lipid binding. Antibodies capable of interacting with the membrane embedded TMDs must also interact with lipids and are thus likely to be polyreactive. Small molecules could in principle either glue or dissociate TMD oligomers to modulate receptor signaling, but the difficulty is also finding the right match between membrane partitioning and TMD interaction. There have been precedence of small molecules targeting receptor TMDs to exert agonistic function. For example, the eltrombopag that activates TpoR by targeting the TMD has been approved for the treatment of idiopathic immune thrombocytopenia^5,41^. In another study of the p75^NTR^, a small molecule was found that can perturb the TMD conformation of the receptor and trigger apoptotic cell death dependent on p75^NTR^ and JNK activity in neurons and melanoma cells^42^. Directly targeting receptor TMD has been explored previously with the use of TM peptides. A TM peptide that competes with TM heterodimerization between the integrin α and β subunits could shift the equilibrium from the inactive to active state^43^. Furthermore, genetic screen of randomly mutated TMD of the bovine papillomavirus E5 protein identified new TMD sequences that can drive PDGF beta receptor dimerization and activation^44^. Similarly, selective inhibition of the erythropoietin receptor (EpoR) and the T-cell receptor by TM peptides has been demonstrated recently^45,46^. These studies suggest that TM helix interactions could afford specificity within the membrane for potentially modulating defective receptor signaling due to TM mutations.

We demonstrate here that, by acquiring solid understanding of the structural basis by which a TMD mutation modulates the activating or inactivating receptor conformation, it is possible to utilize designed TM peptides to specifically suppress the aberrant signaling caused by the TM mutations. In the case of IL-7R, the characterization of a new TMD dimerization mode that supports ligand-independent intracellular JAK1-JAK1 cross phosphorylation provided a TMD design solution to selectively block ligand-independent but not ligand-dependent IL-7R signaling. With the rapid advancements of gene-based intracellular delivery vehicles for protein drugs such as mRNA and AAV^47,48^, the prospect of clinically addressing disease mutations within the receptor TMDs is no longer just a notional concept.

## METHODS

### Reagents and cell lines

Lipids and detergents (POPC, DH^6^PC) were purchased from Avanti Polar Lipids. The deuterated POPC and deuterated DH^6^PC were purchased from Fbreagents. Stable isotopes for NMR spectroscopy experiments were purchased from Cambridge Isotope Laboratories. Phospho-STAT5 rabbit mAb (#9359), phospho-Jak1 rabbit mAb (#74129), STAT5 rabbit mAb (#94205), Jak1 rabbit mAb (#3344), and phospho-STAT5 rabbit mAb AF647 (#9365) were purchased from Cell Signaling Technology. β-Actin rabbit mAb (#AC026) and goat anti-rabbit IgG H+L (HRP) (#AS014) were purchased from ABclonal. Pacific blue anti-hIL-7R mAb (#351306), APC anti-h γc mAb (#338608) and APC anti-Flag Antibody (#637308) were purchased from BioLegend. Recombinant murine IL-3 (#213-13) and recombinant human IL-7 (#200-07) were purchased from PeproTech. Dual-Glo luciferase reporter assay kit (#E2904) was purchased from Promega.

Human embryonic kidney 293T (HEK293T) cells were cultured in Dulbecco’s modified Eagle’s medium (DMEM) supplemented with 10% fetal bovine serum (FBS) and 100 U/mL of pen–strep at 37 °C, 5% CO2. Retroviruses and lentiviruses were produced in HEK293T cells. HEK293T cells were kindly provided by Cell Bank, Chinese Academy of Sciences. Cytokine-dependent BaF3 cells were maintained in RPMI 1640 medium supplemented with 10% FBS and 10 ng/mL of mouse interleukin-3 (mIL-3) at 37 °C, 5% CO2. Phosphorylation of STAT5 and JAK1 were detected in BaF3 cells. HeLa cells were cultured in MEM with Earle’s salts supplemented with 10% FBS superior, 2 mM L-Alanyl-L-Glutamine (Biochrom), 1% non-essential amino acids (Merck KGaA) and 10 mM HEPES buffer (Carl Roth) at 37 °C, 5% CO_2_. Single-molecule imaging and phospho-flow cytometry was performed in HeLa cells.

### Expression and purification of hIL7tm-V253G

The transmembrane domain (TMD) of human IL-7R-V253G (IL-7Rtm-V253G, residues 236-266) was synthesized by GenScript and codon-optimized for E. coli expression. For this construct, C261 was mutated to histidine based on sequence alignment across species, to improve expression and facilitate purification (**Fig. S3**). The expression construct was generated by fusing the IL-7Rtm-V253G fragment to the C terminus of the His9-TrpLE expression sequence in the pMM-LR6 vector (a gift from S. C. Blacklow, Harvard Medical School), with an added methionine between IL-7Rtm-V253G and His9-TrpLE for cleavage by cyanogen bromide (CNBr). The TrpLE tag enables highly hydrophobic and potentially cytotoxic TMDs to be directed into inclusion bodies, thereby increasing expression yields.

For protein expression, transformed *E. coli* BL21 (DE3) cells were grown in M9 minimal medium (50 mM Na_2_HPO_4_, 25 mM KH_2_PO_4_, 10 mM NaCl, 2 mM MgSO_4_, 0.1 mM CaCl_2_, 4 mg/mL Glucose, 1 mg/mL NH_4_Cl and 50 μg/mL kanamycin sulfate) supplemented with Centrum multivitamins and stable isotopes (^15^N, ^13^C) according to the specific requirement for each experiment. Cultures were grown at 37 ℃ to an optical density at 600 nm (OD_600_) of 0.8, then cooled to 25 ℃, and induced with 0.5 mM isopropyl β-D-thiogalactopyranoside (IPTG) overnight at 25 °C. For fully deuterated proteins, bacterial cultures were grown in 99.8% D_2_O (Sigma Aldrich, St. Louis, MO, USA) with deuterated glucose (Cambridge Isotope Laboratories, Tewksbury, MA, USA).

For protein purification, cells were harvested, resuspended in lysis buffer (50 mM Tris, pH 8.0, 200 mM NaCl) and sonicated on ice for 30 min using cycles of 3 s on and 2 s off at 300 W. Inclusion bodies containing the expressed fusion proteins were collected by centrifugation at 18,000 rpm for 30 min, and then resuspended in guanidine buffer (50 mM Tris, pH 8.0, 200 mM NaCl, 6 M guanidine–HCl, and 1% (v/v) Triton X-100) and dissolved overnight at 4 ℃. The fusion proteins were purified by nickel affinity chromatography, followed by CNBr cleavage to remove the His9-TrpLE tag. IL-7Rtm-V253G were further separated by reverse-phase HPLC using a semi-preparative ZORBAX SB-C18 column (Agilent Technologies, Santa Clara, CA, USA) with a 60 min gradient elution from 5–100% buffer B (buffer A: 5% (v/v) isopropanol, 95% H_2_O, 0.1% (v/v) trifluoroacetic acid (TFA); buffer B: 30% (v/v) acetonitrile, 70% isopropanol, 0.1% (v/v) TFA). The fractions containing pure IL-7Rtm-V253G were collected, lyophilized and identified by MALDI-TOF mass spectrometry and SDS-PAGE analysis (**Fig. S2**).

### NMR sample preparation in bicelles

To reconstitute IL-7Rtm-V253G proteins in bicelles, 1–2 mg of purified and lyophilized protein was mixed with 10 mg of protonated or deuterated POPC (Avanti Polar Lipids) and dissolved in hexafluoro-isopropanol (HFIP). The mixture was slowly dried to a thin film under a nitrogen stream and then lyophilized overnight. The dried film was redissolved in 0.5 mL of 25 mM MES buffer (pH 6.5) containing 60 mM protonated or deuterated DH_6_PC (Avanti Polar Lipids). The POPC:DH_6_PC ratio (*q* value) was monitored and verified with 1D ^1^H-NMR. The final NMR samples contained 0.4 to 0.5 mM IL-7Rtm-V253G, ∼30 mM POPC, ∼60 mM DH_6_PC, 25 mM MES (pH 6.5), and 10% D_2_O. For all NOE experiments, proteins were reconstituted in bicelles using POPC and DH_6_PC with deuterated acyl chains (Avanti Polar Lipids).

### NMR backbone resonance assignment

A 0.6 mM (^15^N, ^13^C, 85% ^2^H)-labeled IL-7Rtm-V253G sample in POPC-DH_6_PC bicelles (*q* = 0.5) was used to determine the sequence-specific assignment of backbone amide resonances. The NMR spectra were recorded at 306 K on a Bruker spectrometer operating at ^1^H frequency of 600 MHz with a cryogenic probe. Backbone ^1^HN, ^15^N, ^13^Cα and ^13^C′ resonance assignments were accomplished using the TROSY-based triple resonance experiments, including HNCO, HN(CA)CO, HNCA, HN(CO)CA spectra.

Protein aliphatic and aromatic resonances were assigned using a combination of 2D ^13^C-HSQC, 3D ^15^N-edited NOESY-TROSY-HSQC (mixing time = 120 ms) and ^13^C-edited NOESY-HSQC (mixing time = 150 ms) recorded on a 900 MHz spectrometer. These experiments were performed using a (^15^N, ^13^C)-labeled protein sample reconstituted in deuterated POPC/DH_6_PC bicelles (the acyl chains of POPC and DH_6_PC were fully deuterated). The aliphatic proton resonances were assigned by comparing NOE patterns (specific to α-helical structure) in ^15^N-edited and ^13^C-edited NOE strips. Assignments of the backbone NH and sidechain methyl groups are shown in **Fig. S4**. All the NMR spectra were processed using NMRPipe^49^, and analyzed using XEASY^50^ and CcpNmr^51^.

### Assignment of NOE restraints

For intermolecular distance restraints, we prepared a mixed sample in which half of the protomers were (^15^N, ^2^H)-labeled (0.4 mM) and the other half ^13^C-labeled (0.4 mM), allowing detection of NOEs exclusively between the ^15^N-attached protons of one protomer and the aliphatic protons of the neighboring chain using a 3D ^15^N-edited NOESY-TROSY-HSQC (mixing time = 250 ms). These complementary and reciprocal interchain NOEs identified the helix–helix packing interface.

In addition to interchain NOEs between amide and methyl protons, we recorded another set of 3D ^15^N-edited NOESY-TROSY (mixing time = 120 ms) and 3D ^13^C-edited NOESY-HSQC (mixing time = 150 ms) at 900 MHz using uniformly (^15^N, ^13^C)-labeled IL-7Rtm-V253G. This NOE spectra were used to assign both local and interchain NOEs involving aliphatic protons. The acyl chains of POPC and DH_6_PC used in these samples were also completely deuterated.

### Structure Calculation

The assigned backbone chemical shift values (^15^N, ^1^HN, ^13^Ca, and ^13^C′) from the bicelle-reconstituted protein were used as input for TALOS + program to predict backbone dihedral angles, calculate secondary Cα shifts, and derive the secondary structure of IL-7Rtm-V253G. Backbone dihedral angle restraints (φ = −60°, ψ = −40°) derived from the “GOOD” dihedral angles from TALOS+ were used to define helical regions. All helical regions were restrained using a flat-well harmonic potential (± the corresponding uncertainties from TALOS+, it is ±10°) with the force constant ramped from 5 to 1000 kcal·mol^-1^·rad^-2^.

Structure calculations were performed using XPLOR-NIH^52^. First, the monomer structure of IL-7Rtm-V253G was generated using TALOS+-derived backbone dihedral restraints^53^. Second, the monomer structure and interchain NOEs between amide and methyl protons were used to generate initial homodimer models. Finally, the initial dimer structures were fed to the XPLOR-NIH for iterative refinement against all NMR restraints, while assigning more self-consistent inter- and intrachain NOEs in both ^13^C-edited NOESY-HSQC and isotopically mixed NOE spectra after each iteration. This iterative process resulted in approximately five NOE restraints per residue for the structured regions of the two homodimer complexes (**Table S1**).

The XPLOR refinement used a simulated annealing (SA) protocol in which the temperature in the bath was cooled from 1000 to 200 K with steps of 20 K. The NOE restraints were enforced by flat-well harmonic potentials, with the force constant ramped from 2–30 kcal·mol^-1^·Å^-2^ during annealing. A total of 100 structures were calculated and the 15 lowest-energy structures were selected as the final structural ensemble (**Fig. 1D**). The final ensemble of structures that satisfy all NMR restraints converged to root mean square deviation (RMSD) of 0.631 Å (backbone) and 1.246 Å (heavy atoms) **(Table S1)**.

### Molecular Modeling of IL-7R TMD WT Structure

The wild-type IL-7R transmembrane domain homodimer (IL-7Rtm-WT, residues 235-273; sequence: QGEKDPILLTISILSFFSVALLVILAHVLWKKRIKPVVW) was computationally modeled as a symmetric homodimer using Boltz-2, that is a deep-learning based structural biology foundation model (WT-L255 system)^54,55^. The prediction algorithm was run with the MSA server enabled^56^, pre-trained potentials activated, 50 diffusion samples, and a step scale of 1.0 to ensure adequate conformational sampling and structural diversity in the generated ensemble. Experimentally derived distance constraints from intermolecular NOE cross-peaks (**Fig. 1B**) were incorporated to guide structures toward the observed L255 homodimerization interface. Three constraint sets were implemented reflecting NOE intensity: L255-L255 (strongest NOE, contact restraint ≤ 8.0 Å and bond restraint between methylpropyl carbons), L259-L259 (medium NOE, contact restraint ≤ 10.0 Å), and S252-S252 (weaker NOE, contact restraint ≤ 10.0 Å), with a potential applied to all contact restraints (force: true). From the 50 generated structures, the top 10 models ranked by confidence score were selected for subsequent atomistic molecular dynamics simulations. Model confidence scores are provided in **Fig. S5D**.

### Molecular Dynamics Simulation System Construction

The V253G mutant system (Mutant-G253) was built from the experimentally determined NMR structure with additional N- and C-terminal residues (residues Q235 and 267-273, final sequence: QGEKDPILLTISILSFFSGALLVILAHVLWKKRIKPVVW) added using Modeller^57^ to facilitate proper membrane embedding. Wild-type simulations were performed using two distinct starting configurations to probe both homodimerization interfaces. The first set of WT simulations was generated by back-mutating the V253G NMR structure to the wild-type sequence, corresponding to the G253 interface (WT-V253). The second set of WT simulations utilized the top 10 Boltz-2 predicted structures described above, corresponding to the major L255 homodimerization interface (WT-L255).

All atomistic MD simulation systems (**Table S2**) were constructed using CHARMM-GUI Membrane Builder^58^. Each transmembrane domain homodimer was embedded in a POPC lipid bilayer with explicit TIP3P water molecules and neutralizing Na^+^ and Cl^-^ ions at physiological concentration (150 mM). For all simulation systems, the N-terminus was acetylated and the C-terminus was amidated. All systems were constructed as triclinic boxes with dimensions of approximately 100 × 100 Å in the x-y (membrane) plane (∼140 lipids per leaflet) and 22.5 Å water thickness in the z-direction. All systems followed the CHARMM-GUI six-step equilibration protocol, followed by an additional 3 ns of equilibration with gradually decreasing position restraints on the C_α_ atoms of the transmembrane helix segment (residues 241-264) before production runs. Standard production simulations (**Table S2**) consisted of unbiased MD simulations at 310 K and 1 bar for both WT and V253G systems with durations of 1-2 μs (10 independent repeats).

To assess the relative strength of the dimerization interfaces, temperature scan simulations were conducted where the system temperature was gradually increased from 310 K to 350 K over 200 ns and then kept constant as 350 K for an additional 100 ns to monitor dimer dissociation. A complete summary of all simulation systems and conditions is provided in Table S2.

### Molecular Dynamics Simulation Protocol

All MD simulations were performed using GROMACS 2024^59^ with the CHARMM36m force field^60^ for proteins, CHARMM36 for lipids, TIP3P water model, and standard CHARMM ion parameters under periodic boundary conditions in three dimensions. A 2 fs integration timestep was used with bonds to hydrogen atoms constrained using the LINCS algorithm. Electrostatic interactions were calculated using the smooth particle mesh Ewald method with a real-space cutoff of 1.2 nm. Van der Waals interactions were calculated using a force-switch function between 1.0 and 1.2 nm. Production simulations were conducted in the NPT ensemble at 310 K using the V-rescale thermostat with a coupling constant of 1.0 ps applied separately to membrane, protein, and solvent. Pressure was maintained at 1 bar using the C-rescale barostat with semi-isotropic coupling, a coupling constant of 5.0 ps, and a compressibility of 4.5 × 10⁻⁵ bar⁻¹. For thermal stability simulations, the temperature was linearly increased from 310 K to 350 K over 200 ns after which temperature was kept constant at 350 for another 100 ns.

### Analysis of MD simulations

Trajectory analyses were conducted using custom Python scripts leveraging MDAnalysis^61^ for trajectory handling and Scikit-learn^62^ for clustering, with VMD and PyMOL employed for visualization.

*Thermostability Analysis:* To compare the relative stability of different homodimerization interfaces, we performed temperature scan simulations. The temperature at which each dimer dissociated was used as a measure of interface strength. Dimer dissociation was monitored by tracking the center-of-mass distance between the two transmembrane helices. Dissociation was defined as the point where the helix separation exceeded 4 nm. For each system, observed dissociation temperatures were extracted from multiple independent trajectories.

Dissociation kinetics were modeled using first-order Arrhenius behavior under linear heating following the theoretical framework underlying the Kissinger method for thermally stimulated processes^63^. We assume that (i) the temperature-dependent rate constant follows the Arrhenius behavior, 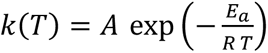, where 𝐴 is the pre-exponential factor and 𝐸*_a_* is the activation energy, and (ii) the loss rate of intact dimers is given by the first-order dissociation kinetics, 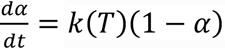, where 𝛼 is the fraction of dissociated dimers. Changing the variable of differentiation in the rate equation from time to temperature using a constant heating rate 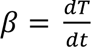, and integrating it from the initial temperature 𝑇_0_ to *T* yields the survival probability, 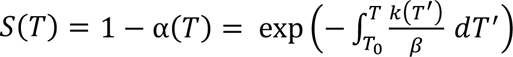, which represents the fraction of dimers remaining intact at temperature 𝑇. Differentiating 𝑆(𝑇) with respect to 𝑇 gives the dissociation probability density per unit temperature, 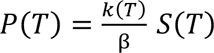.

For each variant, observed dissociation temperatures from multiple independent heating trajectories were used to construct a negative log-likelihood function, which was then minimized to determine the most probable parameter values. 𝑃(𝑇) was used for observed dissociation events, and 𝑆(𝑇) for trajectories where helices had not dissociated at 350 K at the end of the simulation. With only one heating rate, it is not possible to reliably fit both the pre-exponential factor 𝐴 and activation energies 𝐸*_a_* for each variant. Therefore, 𝐴 was treated as a global parameter shared across all variants, while 𝐸*_a_* were fitted individually for each variant. The dissociation temperatures for each variant were then extracted as the 50% survival temperature. Bootstrap resampling was used to estimate uncertainties in the fitted parameters and the dissociation temperatures.

*Clustering Analysis:* To identify distinct conformational states, clustering analysis was performed based on pairwise intermolecular distances between all residue pairs in the two transmembrane helices. For each trajectory frame, the minimum heavy atom distance was calculated for all interchain residue pairs. Frames with minimum interchain distance >5 Å were classified as dissociated and were excluded from further clustering.

For non-dissociated frames, dimensionality reduction was performed using principal component analysis (PCA) on the distance matrix, retaining the first two principal components (**Fig. S5**). A Gaussian Mixture Model (GMM) with two components was fitted to the reduced dataset. Two clusters were sufficient to capture the conformational landscape, as evidenced by clear visual separation in the PCA space and substantially lower within-cluster RMSD compared to between-cluster RMSD (**Fig. S5**).

For each cluster, the centroid structure (frame closest to the Gaussian center) was identified as the representative conformation. Cluster populations for each system were determined as the fraction of frames assigned to each conformational state. Contact maps for each cluster were computed from all frames assigned to that cluster. A residue pair was considered in contact if any heavy atom pair between the two chains was within 4 Å. Contact frequencies were calculated as the fraction of frames in which each residue pair was in contact, yielding an average contact map representative of each conformational state.

### Bacterial adenylate cyclase two-hybrid (BACTH) assay for analysis of TMD interactions

The Bacterial Adenylate Cyclase Two-Hybrid (BACTH) system is based on the interaction-mediated reconstitution of adenylate cyclase (CyaA) activity in *E. coli* and has been used to assess interactions between transmembrane domains (TMDs) in the cell membrane^31^. The BACTH kit (Euromedex, EUK001) was originally purchased from Euromedex and kindly provided by Dr. A. Zoued and R. Ian at Harvard Medical School.

To detect TMDs interaction, the two complementary domains of the CyaA (T18 and T25) were separately fused to the C-termini of IL-7R-TMD or its mutants. An OmpA signal peptide was fused to the N-termini of the TMDs to ensure membrane localization (OmpA-TMD-T18, ampicillin resistance; OmpA-TMD-T25, kanamycin resistance). Both plasmids were co-transformed into the CyaA-deficient *E. coli* strain BTH101. Colonies were selected on LB agar plates containing 50 μg/mL kanamycin and 100 μg/mL ampicillin after incubation at 30 °C for 48 h. Three colonies from each transformation were selected and cultured at 30 °C overnight with shaking (220 rpm) in 3 mL of LB medium supplemented with 100 μg/mL ampicillin, 50 μg/mL kanamycin and 0.5 mM IPTG. Next, 6 μL of each culture was added to the LB-X-Gal reporter plates containing 40 μg/mL X-Gal, 100 μg/mL ampicillin, 50 μg/mL kanamycin, and 0.5 mM IPTG, followed by incubation at 30 °C for 24 h. If no TMD interaction occurs, colonies remain colorless on reporter plates. If TMD interaction occurs, T25 and T18 fragments are brought into close proximity to reconstitute CyaA activity, resulting in blue colonies on reporter plates.

The BolA-like protein IbaG fused to T18 and T25 (IbaG-T18/IbaG-T25) served as a positive control. while the empty vector served as a negative control. Colony color intensity was quantified using ImageJ, and values were normalized to the IbaG-T18/IbaG-T25 positive control. Results are from at least three independent experiments and are expressed as mean ± SEM.

### TOXGREEN assay for analysis of TMD interactions

The TOXGREEN assay is based on the expression of super-folder Green Fluorescent Protein (sfGFP) reporter gene, which depends on the dimerization of the *Vibrio cholerae* transcription factor ToxR. This system has been used to study TMD interactions in the cell membrane^32^, with fluorescent signals detected directly in unprocessed cell cultures. In this system, ToxR and maltose-binding protein (MBP) are fused to the N- and C-termini of the TMD, respectively. The fusion protein (ToxR-TMD-MBP) is expressed in the *E. coli* inner membrane, with the MBP domain located in the periplasm and the ToxR domain in the cytoplasm.

IL-7R-TMD or its mutants were cloned into the pccGFPKAN plasmid (Addgene plasmid #73649) using Gibson Assembly. Glycophorin A (GpA) was used as a positive control (pccGFP-GpA, Addgene plasmid #73651), while the GpA-G83I mutant was used as a negtive control (pccGFP-GpA-G83I, Addgene plasmid #73650). The empty plasmid (pccGFPKAN) was used as a TMD-lacking control, and its background fluorescence was subtracted from all measurements. IL-7R TMD or its mutant plasmids were transformed into *E. coli* strain MM39 (Addgene bacterial strain #42894). Six colonies for each transformation were selected and cultured in 1 mL of LB medium supplemented with 50 μg/mL kanamycin for 18 h at 37 °C. Two hundred microliters of each culture was transferred to a 96-well black-walled, clear-bottom plate. Fluorescence was measured using a multiwell plate reader (EnVision Multilabel Plate Reader; PerkinElmer) at 485 nm excitation and 512 nm emission. Optical density at 600 nm (OD600) was also recorded for normalization.

Data were first normalized by subtracting the average background of the TMD-lacking control, and then normalized to the GpA positive control. Results are from at least three independent experiments and are expressed as mean ± SEM.

### Retroviral production and transduction

Wild-type (WT) or full-length human IL-7R (hIL-7R) mutants were cloned into the retroviral vector MSCV-EGFP with a C-terminal EGFP fusion. WT human γc (hγc) was cloned into MSCV-mCherry with a C-terminal mCherry fusion. Plasmids carrying single-site mutations were generated by site-directed mutagenesis, and all constructs were confirmed by DNA sequencing.

For retrovirus production, HEK293T cells were seeded in 10 cm dishes and grown to 60–70% confluency. Cells were co-transfected with 10 μg retroviral expression plasmid and 8 μg packaging plasmid (pCL-Eco, Addgene, #12371) using Lipofectamine 3000 (ThermoFisher Scientific, #L3000015) according to the manufacturer’s instructions. Six hours after transfection, the medium was replaced with fresh DMEM containing 10% FBS. At 48 h post-transfection, virus-containing supernatants were collected and filtered through a 0.45 μm sterile filter to remove cells and debris. Retroviruses were concentrated using Retro-X Concentrator (TAKARA, #631456) and resuspended in one-hundredth of the original volume of DMEM according to the manufacturer’s instructions.

To infect BaF3 cells, 5 × 10^5^ cells were pelleted and resuspended in retrovirus-containing supernatants supplemented with 10 μg/mL of polybrene and 10 ng/mL of mIL-3. The virus-cell mixture was centrifuged at 1,000 × g for 90 min and incubated overnight at 37 ℃. The next day, the medium was replaced with complete RPMI 1640 medium containing 10% FBS and 10 ng/mL of mIL-3.

To generate stable cell lines, BaF3 cells were sorted by fluorescence-activated cell sorting (FACS, Sony SH800S) based on EGFP (fused to hIL-7R) and mCherry (fused to hγc) fluorescence. EGFP-and/or mCherry-positive cells were collected into complete RPMI 1640 medium containing 10% FBS and 10 ng/mL of mIL-3, cultured for one week, and subsequently used for functional assays.

### Lentiviral delivery of designed transmembrane peptide (TM_PEP_)

To deliver the designed transmembrane peptide (TM_PEP_) into BaF3 cells, cDNAs encoding the WT TM_PEP_ or mutant variants (V253G, L255Y, V253G/L255Y, V253/S249Y) were cloned into the lentiviral vector pLEX. For lentivirus production, HEK293T cells were seeded in 10 cm dishes and grown to 60–70% confluency. Cells were transfected with 6 μg TM_PEP_ plasmid, 4.5 μg packaging plasmid psPAX2 and 1.5 μg envelop plasmid pMD2.G using polyethylenimine (PEI) according to standard protocols. Viral supernatants were harvested 48 h post-transfection, concentrated with PEG8000, and resuspended in DMEM.

For transduction, 5 × 10^5^ BaF3 cells were pelleted and resuspended in lentiviral supernatants supplemented with 10 μg/mL polybrene and 10 ng/mL mIL-3. The virus-cell mixture was centrifuged at 1,000 × g for 90 min and incubated overnight at 37 ℃. The transduced cells were sorted by FACS using APC-conjugated anti-Flag antibody (BioLegend, #637308) at least one week post-transduction. The stably transduced cells were subsequently used for downstream functional assays to assess IL-7R signaling activity.

After 12 hours of cytokine withdrawal, cells were treated with 50 ng/mL IL-7 or PBS for 2 h and then collected for Western blot analysis. Phosphorylated STAT5 (p-STAT5) levels were analyzed to determine whether the designed TM_PEP_ impaired aberrant gain-of-function (GOF) signaling.

### mRNA–LNP delivery of designed transmembrane peptide (TM_PEP_)

To deliver the designed transmembrane peptide (TM_PEP_) into BaF3 cells, chemically modified mRNAs encoding WT TM_PEP_ or mutants (V253G, L255Y, V253G/L255Y, V253/S249Y) were synthesized by GenScript. Each mRNA contained N1-methylpseudouridine, a 5′ Cap1/ARCA, a signal peptide (SP), a FLAG tag, and a 3′ poly (A) tail (∼100 nucleotides). The mRNAs were encapsulated in lipid nanoparticles (LNPs) composed of the ionizable lipid SM-102, structural lipid DSPC, cholesterol, and a PEGylated lipid (DMG-PEG2000). The combination of these components provided particle stability, efficient mRNA encapsulation, and facilitated intracellular delivery.

BaF3 stable cells co-expressing the IL-7R V253G and γc were then treated with the various TM_PEP_-encoding mRNA–LNPs. After 12 h of cytokine withdrawal, the cells were treated with PBS or 50 ng/mL IL-7 for 2 h and collected for immunoblotting analysis.

### Flow cytometry analysis of IL-7R, γc and IL-7R TM_PEP_ expression

BaF3 cells stably expressing WT or mutant human IL-7R, with or without γc, were collected and washed twice with FACS buffer (PBS containing 2% FBS). Cells were resuspended in 200 μL of FACS buffer and incubated on ice for 30 min with Pacific Blue-conjugated anti-hIL-7R antibody (BioLegend, #351306) and APC-conjugated anti-hγc antibody (BioLegend, #338608). To assess expression of WT TM_PEP_ or mutants (V253G, L255Y, V253G/L255Y, V253/S249Y), BaF3 cells stably expressing these constructs were incubated with APC-conjugated anti-Flag antibody (BioLegend, #637308) under the same conditions. After washing three times with PBS, cells were analyzed using flow cytometry (Sony SH800S). Untransfected BaF3 cells were used as a negative control. Data were analyzed using FlowJo 10.8.1 software.

### Western blot analysis of phosphorylated STAT5 and JAK1

To detect STAT5 and JAK1 phosphorylation, BaF3 cells expressing WT or mutant IL-7R or co-expressing with γc were washed three times with PBS and resuspended in RPMI1640 medium without mIL-3 and FBS for 5 h. After cytokine withdrawal, the cells were stimulated with or without 50 ng/mL of hIL-7 for 2 hours at 37 ℃. The cells were harvested and lysed in cell lysis buffer (20mM Tris, 150mM NaCl, 1% Triton X-100) supplemented with protease inhibitor cocktail (Roche, #04693132001) and phosphatase inhibitor cocktail (Roche, #4906845001). Lysates were centrifuged at 12,000 rpm for 10 min at 4 ℃, and protein concentrations were measured with a bicinchoninic acid (BCA) assay. The supernatants were mixed with SDS loading buffer, and equal amounts of protein (30 μg) were loaded for SDS-PAGE. Proteins were transferred to polyvinylidene difluoride (PVDF) membranes (Merck Millipore, #IPVH00010) for 90 min at 300 mA.

PVDF membranes were blocked with 5% non-fat milk in TBST buffer (20 mM Tris, 150 mM NaCl, 0.1% Tween 20, pH 7.6) and incubated overnight at 4 ℃ with primary antibodies against Phospho-STAT5 (Tyr694) Rabbit mAb (Cell Signaling Technology, #9359, dilution 1:500), STAT5 (D2O6Y) Rabbit mAb (Cell Signaling Technology, #94205, dilution 1:1000), Phospho-Jak1(Y1034/1035) (D7N4Z) Rabbit mAb (Cell Signaling Technology, #74129, dilution 1:1000), Jak1 (6G4) Rabbit mAb (D44E3) (Cell Signaling Technology, #3344, dilution 1:1000) and β-actin Rabbit mAb (Abclonal, #AC026, dilution 1:100,000). After washing three times with TBST, membranes were incubated with horseradish-peroxidase (HRP)-conjugated Goat anti-Rabbit IgG (H+L) (Abclonal, #AS014) for 60 min at room temperature. protein bands were detected using Pierce ECL Western blotting substrate (ThermoFisher Scientific, #32106) or Immobilon Western HRP Substrate (Millipore, #WBKLS0050) according to the manufacturer’s instruction. The intensity of immunoblot bands was quantified using ImageJ software^64^.

### Luciferase reporter assay to assess JAK–STAT5 signaling pathway activation

To assess IL-7R activation in BaF3 cells stably expressing WT or mutant human IL-7R, the pHAGE-STAT5-luciferase reporter plasmid was introduced via lentiviral transduction. At 48 h post-infection, cells were washed three times with PBS and starved 5 h in RPMI 1640 medium without mIL-3 and FBS. cells were then seeded at 5 × 10^4^ cells per well in white, clear-bottom 96-well cell culture plate. Luminescence was measured using the Dual-Luciferase Reporter Assay System (Promega) according to the manufacturer’s instructions. Data were analyzed using GraphPad Prism 9 software and results are shown as mean ± SEM from three independent experiments (n = 3).

### Flow cytometry analysis of STAT5 phosphorylation (p-STAT5)

To quantify STAT5 phosphorylation activity of tagged IL-7R variants, HeLa cells were transiently transfected with IL-7R (residues 37-459) or IL-7RΔECD (residues 236-459) and pSems-JAK1-mEGFP using polyethyleneimine (PEI) in 6 cm dishes for 5 h at 37 °C. After overnight incubation, cells were detached using Accutase and transferred into 10 mL Falcon tubes. Cells were pelleted by centrifugation at 300 × g for 5 min at 4 °C and resuspended in 200 µL PBS supplemented with 0.5% BSA (PBSA). Cells were transferred into V-bottom 96-well plates and then fixed with 4% PFA for 10 min at room temperature. After fixation, the cells were centrifuged at 300 × g for 5 min at 4 °C, and permeabilized with ice-cold methanol for 30 min on ice. After permeabilization, cells were centrifuged again at 300 × g for 5 min and washed three times with PBSA. Subsequently, the cells were incubated with AlexaFluor 647-conjugated anti-p-STAT5 antibody (Cell Signaling Technology, #4324, 1:100 in PBSA) and 10 nM anti-GFP nanobody-enhancer conjugated to Dy410 for 1 h at room temperature. Cells were washed three times and analyzed by flow cytometry. Live cells positive for IL-7R/IL-7RΔECD and JAK1 expression were gated, and 3,000–6,000 cells per sample were analyzed. Data are presented as median fluorescent intensity (MFI) ± SEM from a representative experiment.

### Single-molecule receptor dimerization assays

For single-molecule imaging, IL-7R (template kindly provided by Shai Izraeli, Tel Aviv University) was cloned into the vector pSems including the signal sequence of Igκ (pSems leader) and a suitable N-terminal tag for cell surface labeling upstream of the receptor. Full-length IL-7R (37-459) was fused to an N-terminal ALFAtag (pSems leader ALFA-IL-7R WT and V253G, respectively) and IL-7RΔECD (236-459) was fused to nonfluorescent monomeric GFP (pSems leader mXFP-IL-7RΔECD WT and V253G, respectively). JAK1 constructs (JAK1ΔTK and JAK1ΔPKTK) were cloned into the vector pSems and C-terminally fused to mEGFP (pSems-JAK1ΔTK-mEGFP and pSems-JAK1ΔPKTK-mEGFP, respectively), which was used to confirm expression before image acquisition. The day before transfection, HeLa cells were detached using trypsin and seeded onto 6 cm dishes. After overnight incubation, cells were transfected with IL-7R (37-459) or IL-7RΔECD (236-459) and pSems-JAK1ΔTK-mEGFP. For transient transfection, plasmid DNA was mixed with polyethylenimine (PEI) in 300 µL 150 mM NaCl for 15 min and then added drop-wise to the 6 cm dishes. After 4.5 h incubation at 37 °C, transfection was stopped by washing three times with PBS and the cells were detached and seeded onto 25 mm glass coverslips coated with a poly-L-lysine-graft-(polyethylene glycol) copolymer functionalized with RGD (PLL-PEG-RGD) to support cell adhesion while minimizing non-specific binding of fluorescent nanobodies.

IL-7R dimerization was quantified by dual-color single-molecule FRET localization and tracking as described previously^20^. IL-7R was labeled via a equimolar mixture of ALFA-tag nanobodies (nbALFA) for full-length constructs, or GFP-enhancer nanobodies (nbEN) for truncated constructs, which were site-specifically conjugated to a fluorophore via a cysteine residue by maleimide chemistry. Imaging experiments were carried out 12-16 h post transfection at 25°C in medium without phenol red and supplemented with an oxygen scavenger and redox-active photoprotectant (0.5 mg mL^−1^ glucose oxidase (Sigma-Aldrich), 0.04 mg mL^−1^ catalase (Sigma-Aldrich), 5% w/v glucose (Carl Roth), 1 μM ascorbic acid (Carl Roth) and 1 μM methylviologen(Sigma-Aldrich)) to minimize photobleaching). For cell surface labeling of ALFA-tagged receptors, a mixture of nbALFA conjugated to Cy3B and ATTO643 (degree of labeling: 1.0), respectively, were added to the medium at equal concentrations (3 nM each) and kept in the medium during the experiment. After 5 min of incubation, image acquisition was initiated while the labeled nanobodies were maintained in the bulk solution throughout the experiment to ensure equilibrium binding. smFRET time-lapse imaging by ALEX was carried out by dual-color total internal reflection fluorescence microscopy using an inverted microscope (IX71, Olympus) equipped with a spectral image splitter (DualView, Optical Insight). Donor and acceptor dyes were sequentially excited at 561 nm and 642 nm, respectively, for 25 ms per channel, resulting in an overall frame rate of 20 frames per second. Before image acquisition, co-expression of JAK1ΔTK or JAK1ΔPKTK was confirmed by mEGFP fluorescence upon excitation at 488 nm. Images were acquired by an EMCCD camera (iXon Ultra, Andor). Alternating laser excitation was achieved with a micro-controller (Arduino Uno) and open-source acquisition software^65^ synchronizing laser shuttering via an AOTF and camera triggering. For each cell, image stacks of 400 consecutive frames were recorded.

### Single-molecule image analyses

Single-molecule image analyses was performed using custom-written MATLAB code (SLIMfast 4C^33^). In brief, cell-specific regions of interest (ROIs) were calibrated across all three imaging channels (donor, acceptor, and FRET) using fluorescent fiducial markers to ensure spatial alignment. Molecule positions were then determined by fitting a fixed point spread function (PSF) model, providing high-precision localizations. To minimize background and artifacts, immobile emitters were excluded by spatial cluster filtering^35^. In addition, donor and acceptor trajectories shorter than 10 consecutive frames (500 ms) were discarded, as these likely represent random events rather than true molecular signals. The filtered trajectories were then converted back into localization data, which enabled robust co-localization with FRET signals. This step was critical to reduce false-positive events arising from spectral crosstalk and thus ensured a reliable identification of specific FRET interactions. Based on this data, the relative fraction of dimerized receptors (𝑓^345^) was calculated from the overlap of filtered donor and acceptor localizations (𝑛^2∗^ and 𝑛^7∗^) with co-localized smFRET events (𝑛^87∗^). To correct for variations in dual-color labeling efficiency, including dimers labeled with the same fluorophore and the labeling ratio across both channels (𝑐_5_), the following correction was applied:

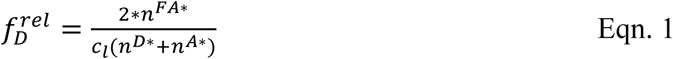

This approach provides an accurate quantification of receptor dimerization by correcting for labeling bias while excluding spurious co-localizations. To further investigate the stability of dimers, we tracked molecules localized in the FRET channel. Here, immobile or noisy signals were removed by cluster filtering, and trajectories were only considered if their diffusion coefficient was below 1.5 µm²/s, thus ensuring exclusion of background fluorescence. The resulting trajectory lengths were compiled into histograms and fitted with an exponential decay model, allowing the extraction of dimer lifetimes.

The diffusion properties of monomers and dimers were determined from pooled single-molecule trajectories using mean squared displacement (MSD) analysis in MATLAB. The resulting MSD values were converted into diffusion coefficients^33^:

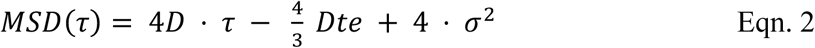

Where τ is the lag time, D the diffusion coefficient, 𝑡𝑒 the exposure time, and σ the localization precision. For monomers, only trajectories exceeding 10 frames were included, whereas for dimers the analysis was restricted to FRET trajectories with more than 5 frames and a half-time threshold above 300 ms per cell.

FRET efficiencies were quantified using a recently published custom-written MATLAB (MathWorks, R2022b) script^66^, considering the fluorescence signals from the three relevant channels of ALEX experiments: the directly excited donor, the directly excited acceptor, and the sensitized FRET channel. Initially, the signals from the directly excited donor and the FRET channel are combined to form a merged donor-FRET signal, which is then co-localized with the directly excited acceptor signal. The apparent FRET efficiency *E_raw_* was calculated for each pair of co-localized emitters according to:

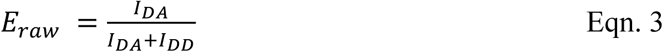

where *I_DA_* and *I_DD_* represent the integrated single emitter intensities in the FRET and donor channels, respectively. To obtain accurate FRET efficiencies *E_corr_*, signals were corrected for background and bleed-through, and the microscope and FRET-pair dependent semi-empiric γ-factor of 0.45 was applied^66^. Histograms generated from all *E_corr_* values obtained from multiple cells were fitted by multiple Gaussian distributions, accounting for random co-localizations (*E_o_* ≈ 0) and 2-3 additional peaks.

### Statistical analysis

GraphPad PRISM v9 and Origin v9 were used for statistical analyses. Unpaired two-sided Student’s t-test was performed to compare two groups of independent samples. One-way analysis of variance (ANOVA) was performed to compare three or more groups of independent samples. Western blot and BACTH results were based on three independent experiments and analyzed with ImageJ v1.53n. Flow cytometry results were based on two or three independent experiments and analyzed with FlowJo v10. *p* < 0.05 was considered statistically significant. Additional information can be found in the figure legends.

### Data availability

The atomic structure coordinate and structural constraints have been deposited in the Protein Data Bank (PDB) with accession numbers 9XEA (Structure of the transmembrane domain dimer of human IL-7R V253G mutant). ^1^H, ^13^C, and ^15^N chemical shift values have been deposited in the Biological Magnetic Resonance Data Bank (BMRB) with accession numbers 36805.

## Acknowledgements

We thank A. Zoued and R. Ian for sharing materials for the BACTH assay, the staff at the Nuclear Magnetic Resonance System/Mass Spectrometry System/Integrated Laser Microscopy System at the National Facility for Protein Science in Shanghai for technical support and assistance in data collection and analysis, and A. Budke-Gieseking, G. Hikade, and W. Kohl for technical support. We thank CSC – IT Center for Science Ltd (Espoo, Finland) for providing computing resources to perform molecular dynamics simulations. We thank Shreyas Kaptan and Felix Eurasto for their help with clustering.

## Funding

This work was supported by Science and Technology Commission of Shanghai Municipality (22JC1410500), the National Natural Science Foundation of China-Research Fund for International Scientists (82350710799), the Strategic Priority Research Program of the Chinese Academy of Sciences (XDB1060000), the Shanghai Basic Research Pioneer Project, the Shanghai Municipal Science and Technology Major Project, the Chinese Academy of Sciences (2023000040, SIOCIRCBC230202) (to J.J.C.), and the Deutsche Forschungsgemeinschaft (DFG, German Research Foundation, SFB 1557, 467522186) (to J.P.), the Research Council of Finland grants (331349, 336234, and 346135), the Sigrid Juselius Foundation, Helsinki Institute of Life Science (HiLIFE) Fellow Program, and the Lundbeck Foundation (to I.V.). It was also supported by the China Postdoctoral Science Foundation (2024M763430) and the Shanghai Postdoctoral Excellence Program (2024729) (to Q.W.).

## Author contributions

T.C., J.J.C., Q.W., and J.P. conceived the study. Q.W., Z.D., and T.C. performed structural analyses of the mutant by NMR and biochemical analyses by BACTH and TOXGREEN. Q.W., A.Z.C., M.C., W.B, and T.C. designed and performed cell signaling assays. M.C., T.C., Q.W., and J.J.C. designed and performed the transmembrane peptide interference studies. A.L. and C.P. designed, performed and analyzed single-molecule imaging experiments. S.V., O.H., G.E., and I.V. designed the biomolecular simulations. G.E. and I.V. supervised the biomolecular simulation work. O.H., S.V., and G.E. built the molecular models. S.V. performed the simulations. G.E. and S.V. analyzed the data. J.J.C., Q.W., and J.P. wrote the paper and all authors contributed to editing of the manuscript.

## Competing interests

The authors declare no competing interests.

## Notes

### Competing Interest Statement

The authors have declared no competing interest.

