## Supplemental Figures and Tables for "Structural Rewiring of IL-7R Dimerization by an Oncogenic Transmembrane Mutation Can Be Reversed by Rational Design"

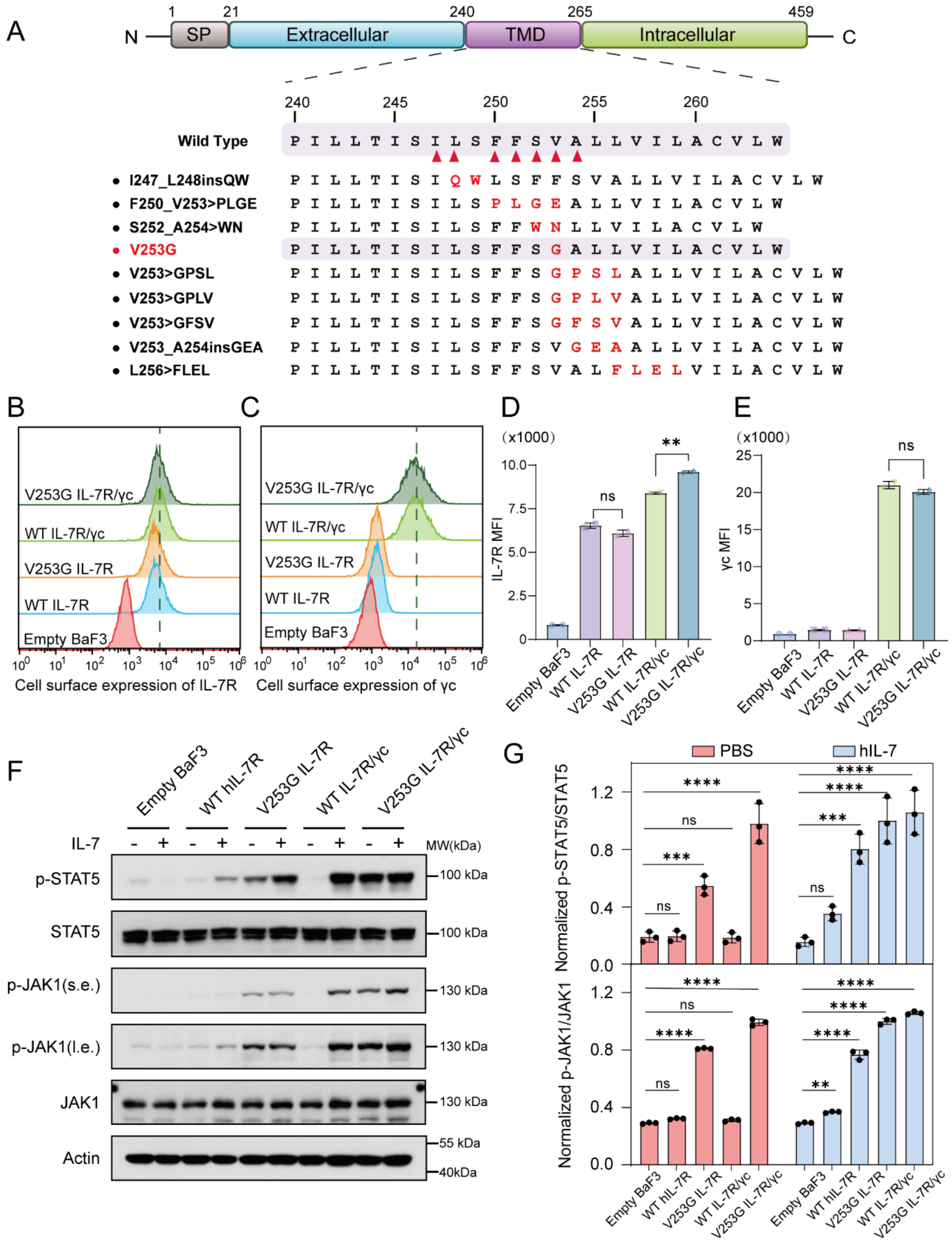


**Fig. S1. V253G mutation causes ligand independent receptor signaling of IL-7R.**

(**A**) Summary of gain-of-function (GOF) mutations in human IL-7R TMD associated with T-ALL. *Top*: domain organization of IL-7R highlighting the TMD (residues 240–264); red triangles indicate mutation sites. *Bottom*: the amino acid sequences of GOF mutants are listed with substituted or inserted residues in red.

(**B**) Flow cytometry analyses of the WT and V253G IL-7R surface expression alone and with co-expression of γc in BaF3 cells, as reported by Pacific Blue-conjugated anti-IL-7R antibody. Untransfected (Empty) BaF3 cells were used as negative controls.

(**C**) The same analyses as in (B) but reported by APC-conjugated anti-γc antibody.

(**D-E**) Quantification of IL-7R and γc surface expression levels using the mean fluorescence intensity (MFI) in (B) and (C), respectively.

(**F**) Phosphorylation of STAT5 (p-STAT5) and JAK1 (p-JAK1) in WT, V253G, WT/γc, and V253G/γc BaF3 cells, with or without treatment with 50 ng/mL IL-7. The p-STAT5, p-JAK1, STAT5 and JAK1 signals were detected by immunoblotting.

(**G**) Quantification of p-STAT5 and p-JAK1 signals in (F) as p-STAT5/STAT5 and p-JAK/JAK1 intensity ratios, normalized to that of the WT IL-7R/γc receptors activated by IL-7. The data are shown as means ± SEMs calculated from three independent experiments. Statistical significance by One-way ANOVA. ns, not significant; *$p \leq 0.05$; ** $p \leq 0.01$; *** $p \leq 0.001$; **** $p \leq 0.0001$. PBS, phosphate-buffered saline.

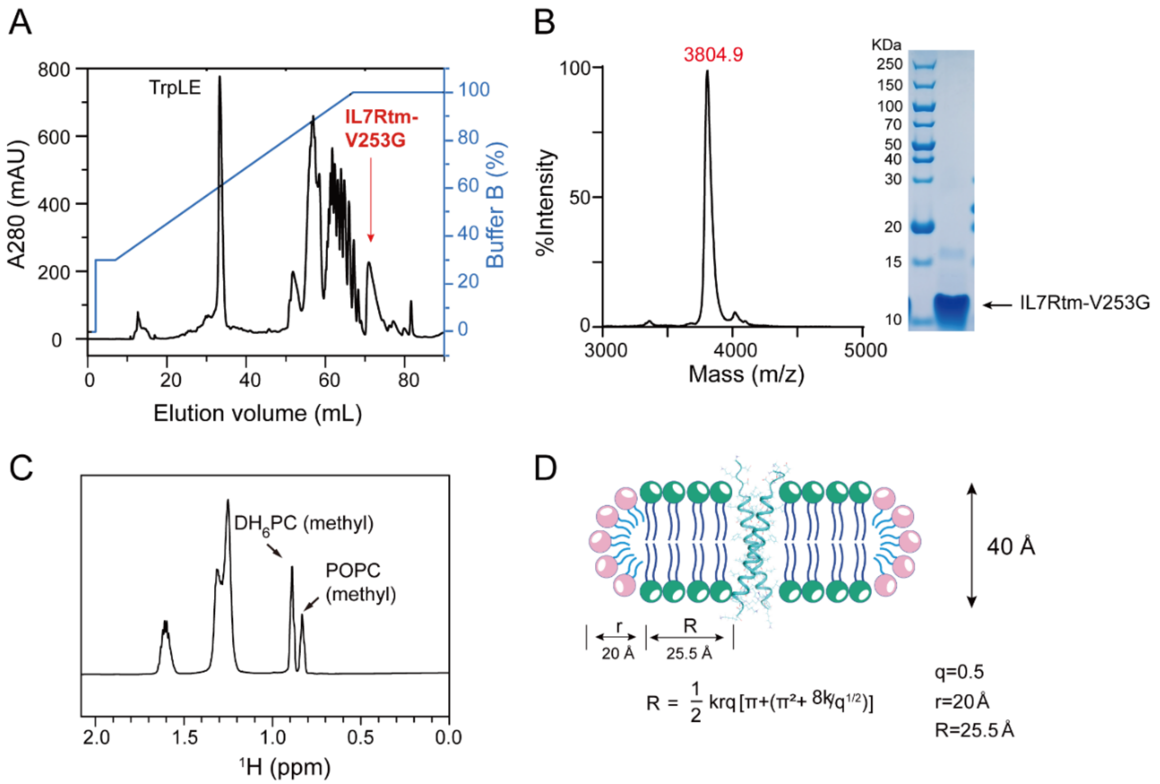


**Fig. S2. Expression, purification, and bicelle reconstitution of IL-7Rtm-V253G.**

(**A**) Reverse phase HPLC purification of human IL-7Rtm-V253G from CNBr-cleaved TrpLE- IL-7Rtm-V253G on a Zorbax SB-C3 column with a gradient from 30-100% acetonitrile (ACN) and 0.1% trifluoroacetic acid (TFA). The purified IL-7Rtm-V253G product was verified with SDS-PAGE and mass spectrometry.

(**B**) MALDI-TOF mass spectrometry signal (left) and SDS-PAGE results (right) showing HPLC-purified IL-7Rtm-V253G protein. The molecular weight (MW) of IL-7Rtm-V253G was measured as 3804.9 Da.

(**C**) $^1$H NMR spectrum of the $DH_6PC$ and POPC methyl peaks from IL-7Rtm-V253G NMR sample. The integrated areas of the $DH_6PC$ and POPC methyl peaks used to calculate the bicelle q value were 0.5.

(**D**) Schematic illustration of IL-7Rtm-V253G reconstituted in POPC/$DH_6PC$ bicelles with the planar lipid bilayer formed by POPC in a radius of 25.5 Å and the micellar lipid rim formed by $DH_6PC$ in a radius of 20 Å.

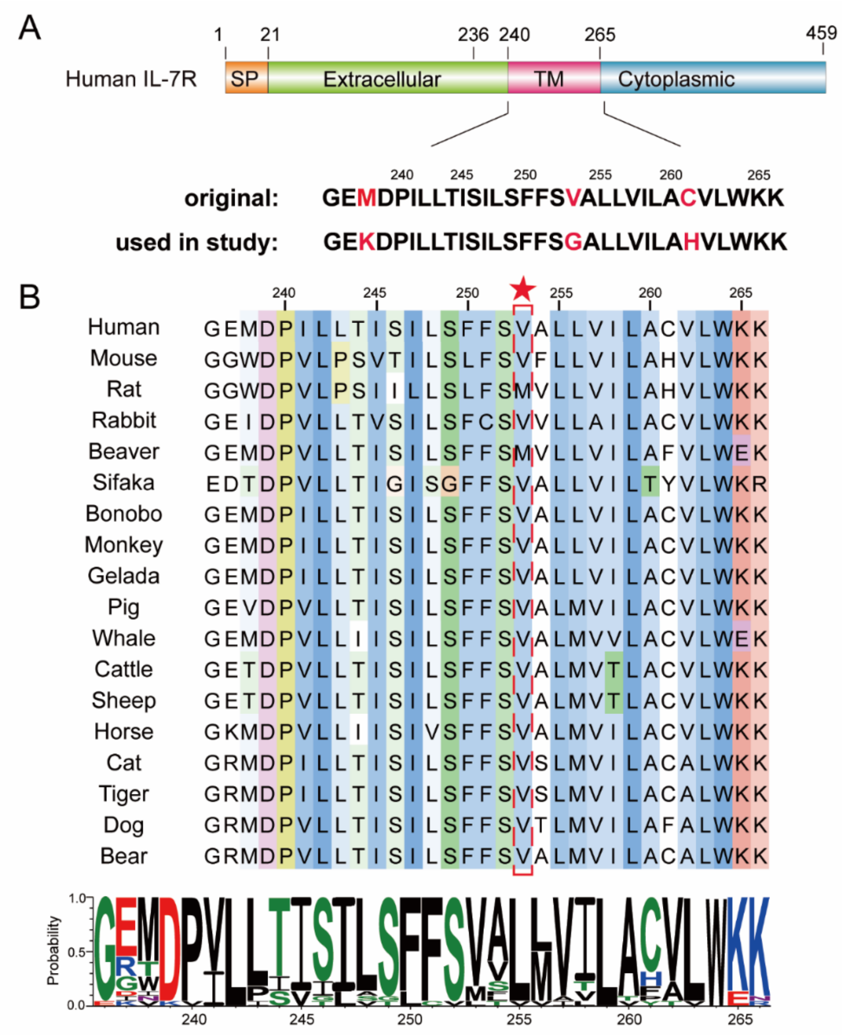


**Fig. S3. Evolutionary conservation of key IL-7R features**

(**A**) The domain organization of human IL-7R (hIL-7R) and the amino acid sequence of hIL-7R TMD (residues 236-266). The hIL-7Rtm-V253G constructs used in this study are shown with the cysteine and methionine residues (colored in red) mutated.

(**B**) Eighteen vertebrate species were chosen for IL-7R protein alignment. The sequence logo (lower panel) illustrates the degree of amino acid conservation based on hIL7R TMD alignments [67]. The relative conservation of each amino acid residue is indicated by the height of each letter stack. Amino acids are color-coded as basic (blue), acidic (red), neutral (purple), polar (green), or hydrophobic (black). We noticed that V253 is highly conserved among these species.

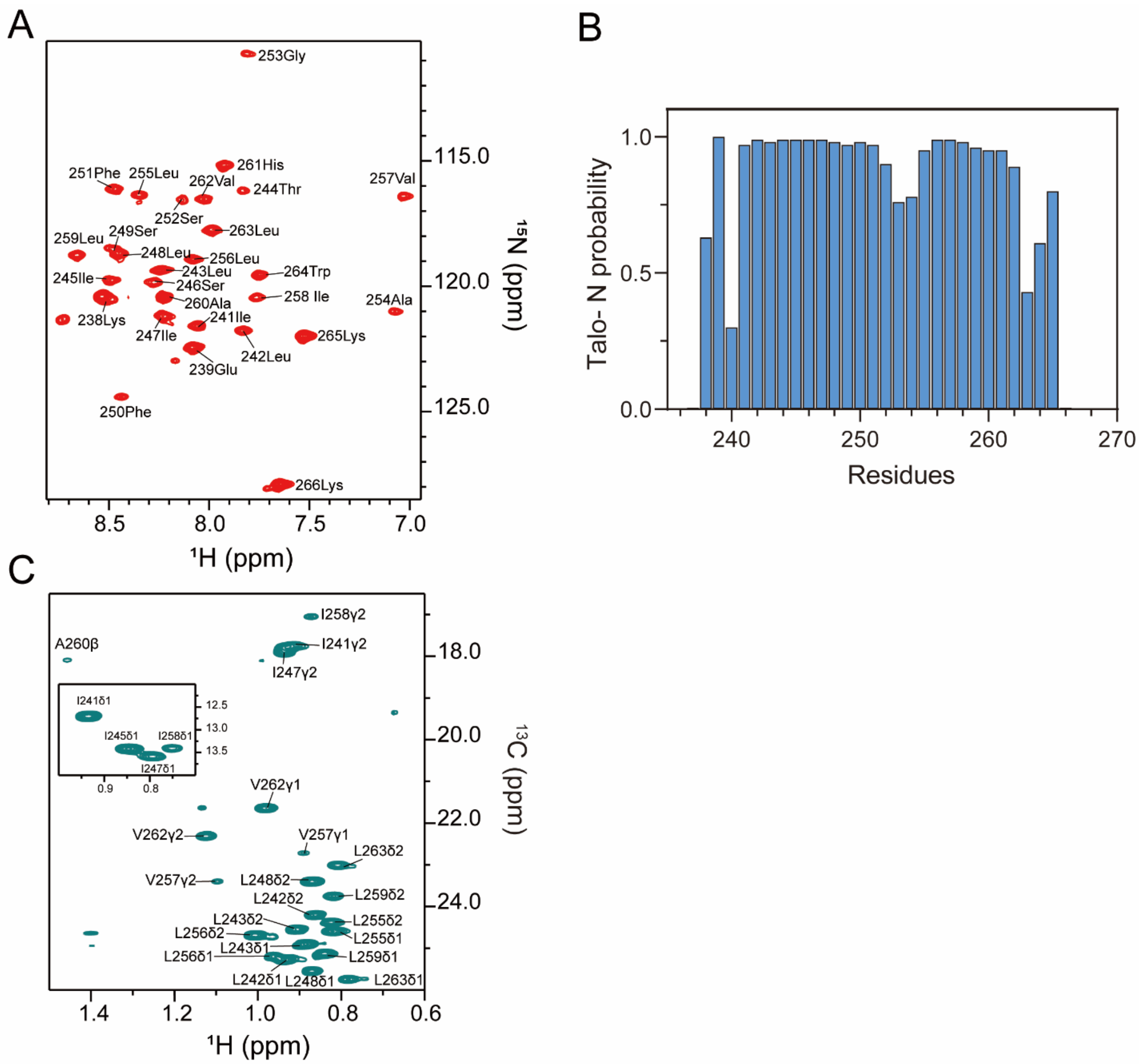
A
253Gly
261His
251Phe
255Leu
262Val
244Thr
257Val
252Ser
263Leu
259Leu
249Ser
248Leu
256Leu
243Leu
264Trp
245Ile
246Ser
258 Ile
260Ala
254Ala
238Lys
241Ile
247Ile
265Lys
242Leu
239Glu
250Phe
266Lys
115.0
120.0
125.0
¹⁵N (ppm)
8.5
8.0
7.5
7.0
¹H (ppm)
B
1.0
0.5
0.0
Talo- N probability
240
250
260
270
Residues
C
I258γ2
I241γ2
A260β
I247γ2
I241δ1
I245δ1
I258δ1
I247δ1
12.5
13.0
13.5
0.9
0.8
18.0
20.0
22.0
24.0
¹³C (ppm)
V262γ1
V262γ2
V257γ1
L263δ2
V257γ2
L248δ2
L259δ2
L242δ2
L243δ2
L255δ2
L256δ2
L255δ1
L243δ1
L256δ1
L259δ1
L242δ1
L248δ1
L263δ1
1.4
1.2
1.0
0.8
0.6
¹H (ppm)

**Fig. S4. NMR characterization of the IL-7Rtm-V253G in bicelles.**

(**A**) The 2D $^{1}H$, $^{15}N$ TROSY-HSQC spectrum of IL-7Rtm-V253G samples in POPC/$DH_6PC$ bicelles recorded at 306 K on a Bruker 600 MHz spectrometer.

(**B**) Chemical-shift-based secondary structure prediction of IL-7Rtm-V253G in bicelles. The graph shows the probability of each residue being part of the α helix, as determined with TALOS+ software [53].

(**C**) The methyl group region of the 2D $^{1}H$,$^{13}C$ HSQC with 28 ms constant-time $^{13}C$ evolution, recorded at an $^{1}H$ frequency of 900 MHz, using ($^{15}N$, $^{13}C$)-labeled IL-7Rtm-V253G protein.

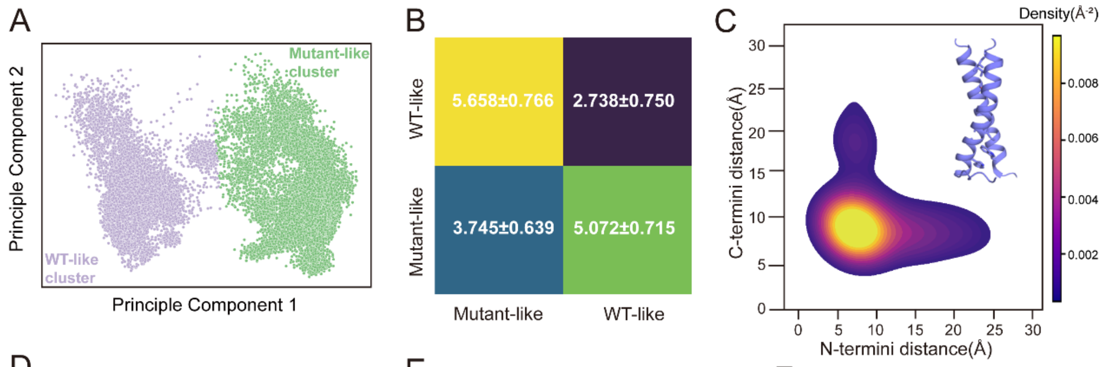


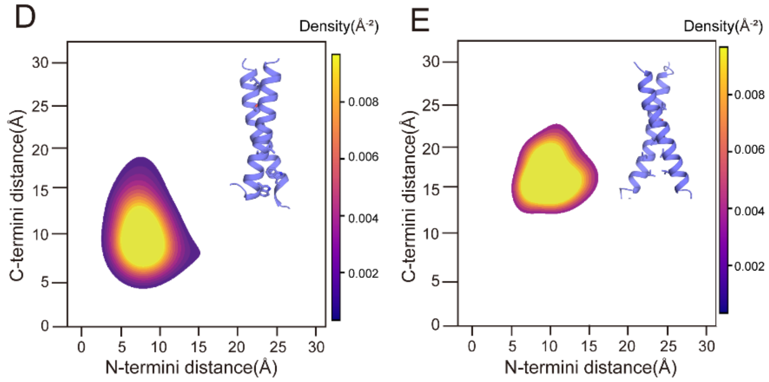


F

| Model | Confidence | pLDDT | ipTM | pDE |
|---|---|---|---|---|
| 0 | 0.699 | 0.733 | 0.559 | 0.640 |
| 1 | 0.695 | 0.738 | 0.523 | 0.609 |
| 2 | 0.694 | 0.742 | 0.506 | 0.673 |
| 3 | 0.693 | 0.725 | 0.568 | 0.634 |
| 4 | 0.691 | 0.733 | 0.527 | 0.690 |
| 5 | 0.686 | 0.726 | 0.529 | 0.681 |
| 6 | 0.683 | 0.729 | 0.500 | 0.710 |
| 7 | 0.682 | 0.728 | 0.502 | 0.640 |
| 8 | 0.679 | 0.726 | 0.493 | 0.679 |
| 9 | 0.679 | 0.730 | 0.473 | 0.695 |

**Fig. S5. Validation of clustering, geometric differences between conformational states, and WT-L255 model quality**

**(A)** PCA projection of non-dissociated frames, colored by Gaussian Mixture Model cluster assignment (Mutant-like Cluster, purple; WT-like Cluster, green). Clear separation validates the two-state conformational landscape.

**(B)** Within-cluster and between-cluster root-mean-square deviations (RMSD ± SE) confirm distinct conformational states with substantially lower within-cluster RMSD compared to between-cluster RMSD.

**(C–E)** Density distributions of the distance between the COM of N-terminal residues 241–242 and C-terminal residues 263–264 for each cluster, illustrating the distinct geometries of the two conformational states. The color bar in panel E applies to all maps.

**(F)** Confidence metrics for the ten highest-scoring Boltz-2 predicted WT-L255 dimers. Confidence represents overall prediction quality, pLDDT indicates per-residue structure confidence (0–1 scale), ipTM measures interface prediction quality, and pDE estimates distance error in units of Å. ipTM > 0.5 indicates reliable interface predictions.

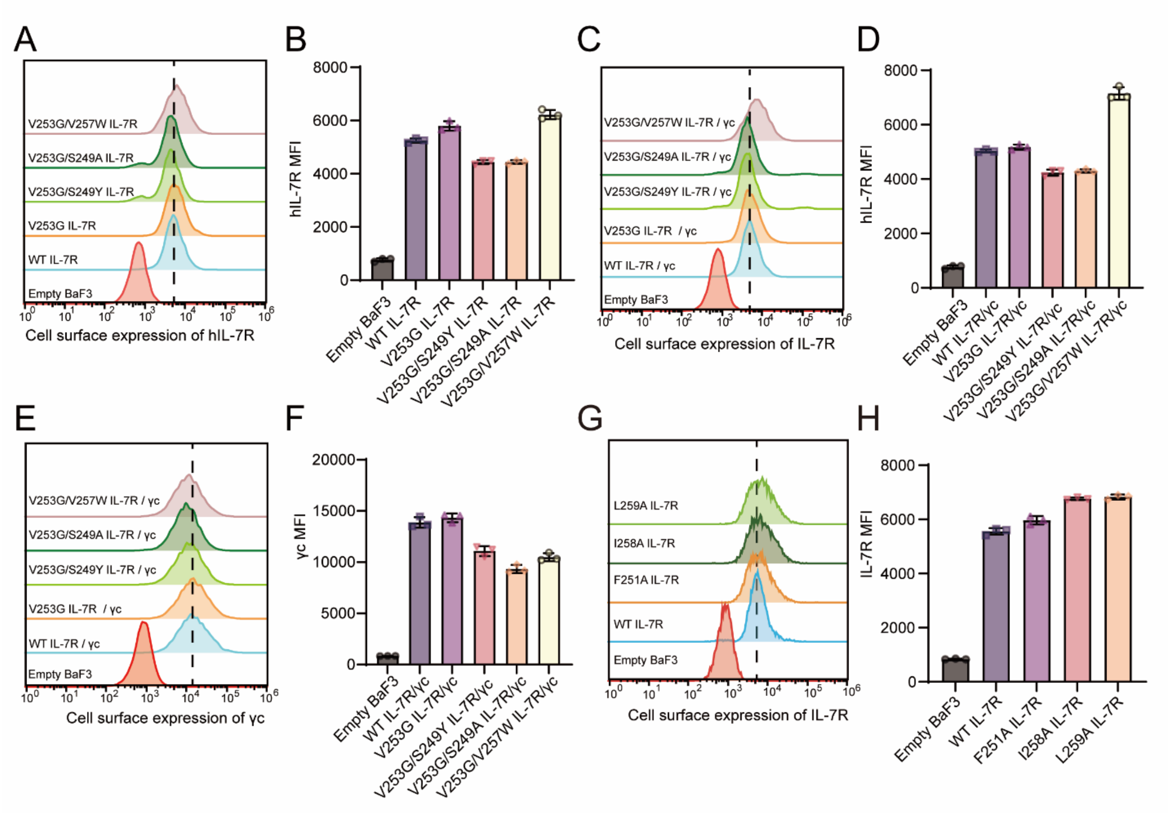


**Fig. S6. Flow cytometry analysis of cell surface coexpression of WT or mutants of IL-7R.**

(**A**) Flow cytometric analysis of IL-7R membrane expression (V253 dimerizing face) in BaF3 cells expressing IL-7R alone. Untransfected (Empty) BaF3 cells were used as negative controls.

(**B**) Geometric mean fluorescence intensity of IL-7R surface expression corresponding to (A).

(**C**) Flow cytometric analysis of IL-7R membrane expression in BaF3 cells co-expressing IL-7R and γc. Untransfected (Empty) BaF3 cells were used as negative controls.

(**D**) Geometric mean fluorescence intensity of IL-7R surface expression corresponding to (C).

(**E**) Flow cytometric analysis of γc membrane expression in BaF3 cells co-expressing IL-7R and γc. Untransfected (Empty) BaF3 cells were used as negative controls.

(**F**) Geometric mean fluorescence intensity of γc surface expression corresponding to (E).

(**G**) Flow cytometric analysis of IL-7R membrane expression (L255 dimerizing face) in BaF3 cells expressing IL-7R alone. Untransfected (Empty) BaF3 cells were used as negative controls.

(**H**) Geometric mean fluorescence intensity of IL-7R surface expression corresponding to (G).

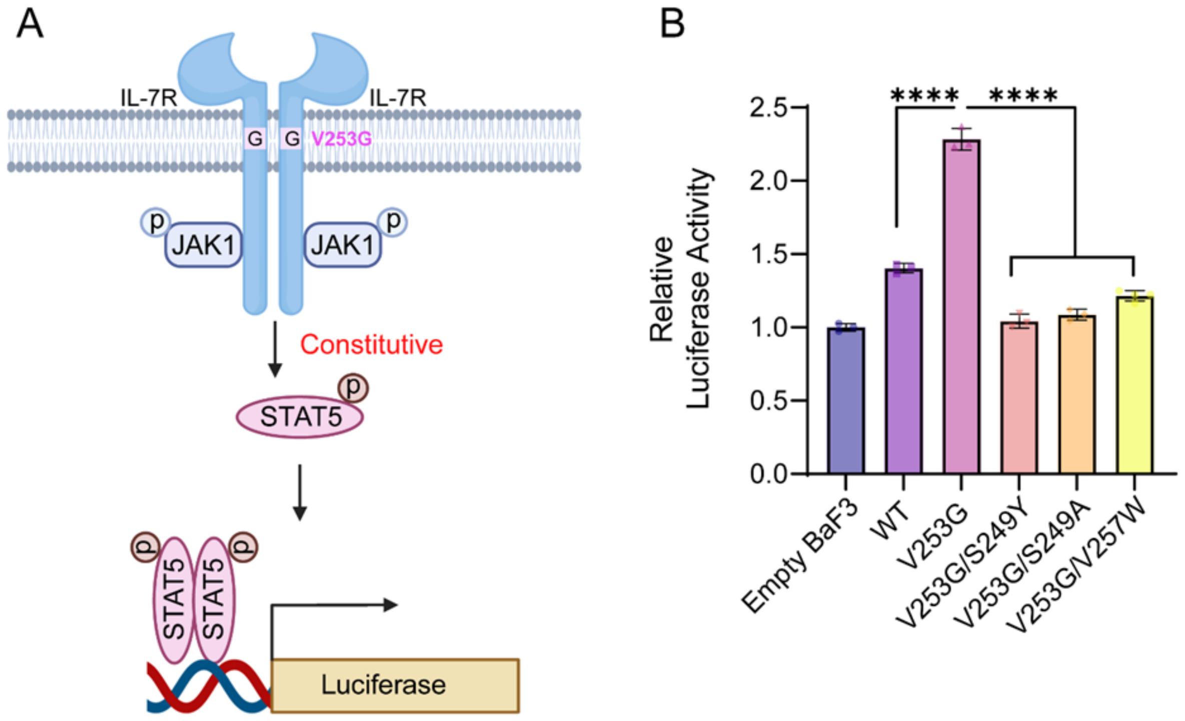


**Fig. S7. Luciferase reporter assay analysis of STAT5 signaling pathway activity of WT or mutants of IL-7R.**

(**A**) Schematic representation of the STAT5 luciferase reporter assay. The luciferase reporter vector containing a STAT5 response element that drives transcription of the luciferase gene. BaF3 cells co- transfected with the reporter plasmid and IL-7R plasmid were used to monitor the STAT5 signaling pathway activity.

(**B**) Luciferase reporter assay analysis of STAT5 transcriptional activity in BaF3 cells expressing wild-type or mutant IL-7R, normalized to Empty BaF3. Data are shown as means ± SEMs calculated from all technical replicates from three independent experiments. Statistical significance by One-way ANOVA. ns, not significant; *$p \leq 0.05$; ** $p \leq 0.01$; *** $p \leq 0.001$; **** $p \leq 0.0001$.

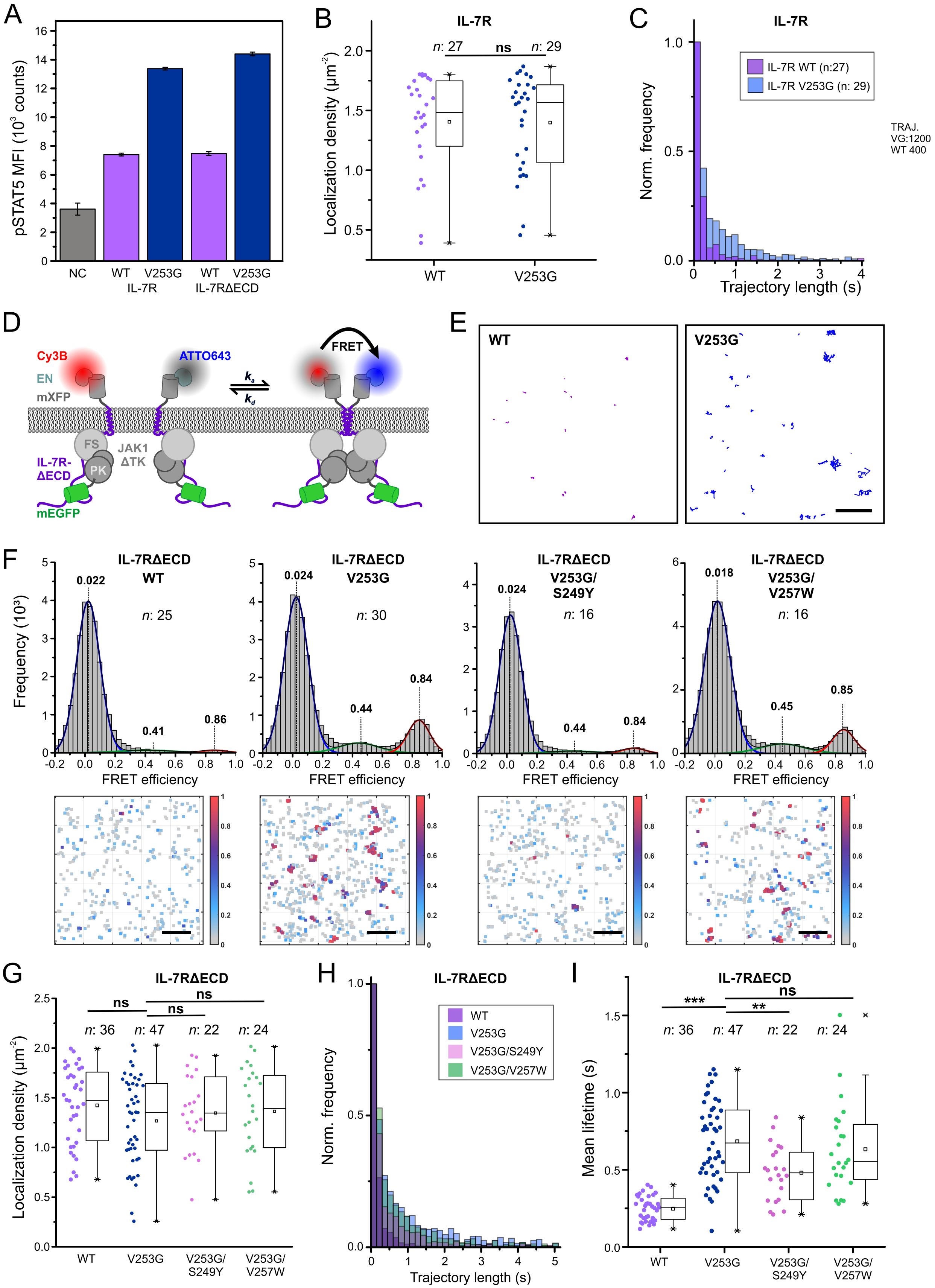

A
pSTAT5 MFI (10³ counts)
NC
WT
V253G
IL-7R
WT
V253G
IL-7RΔECD
B
IL-7R
n: 27
ns
n: 29
Localization density (µm⁻²)
WT
V253G
C
IL-7R
IL-7R WT (n:27)
IL-7R V253G (n: 29)
Norm. frequency
Trajectory length (s)
TRAJ.
VG:1200
WT 400
D
Cy3B
EN
mXFP
ATTO643
FRET
FS
PK
JAK1
ΔTK
IL-7R-
ΔECD
mEGFP
E
WT
V253G
F
IL-7RΔECD
WT
n: 25
IL-7RΔECD
V253G
n: 30
IL-7RΔECD
V253G/
S249Y
n: 16
IL-7RΔECD
V253G/
V257W
n: 16
Frequency (10³)
FRET efficiency
G
IL-7RΔECD
ns
ns
ns
n: 36
n: 47
n: 22
n: 24
Localization density (µm⁻²)
WT
V253G
V253G/
S249Y
V253G/
V257W
H
IL-7RΔECD
WT
V253G
V253G/S249Y
V253G/V257W
Norm. frequency
Trajectory length (s)
I
IL-7RΔECD
***
ns
**
n: 36
n: 47
n: 22
n: 24
Mean lifetime (s)
WT
V253G
V253G/
S249Y
V253G/
V257W

**Fig. S8. Quantifying IL-7R dimerization by smFRET.**

(**A**) p-STAT5 activation determined by phospho-flow cytometry compared for IL-7R and IL-7RΔECD, WT and V253G, respectively. Negative control (NC) in the absence of IL-7R.

(**B**) IL-7R cell surface density observed in different smFRET dimerization experiments.

(**C**) Normalized smFRET trajectory length histograms for IL-7R WT and V253G.

(**D**) Cartoon of smFRET dimerization assays using IL-7RΔECD N-terminally tagged with a mXFP-tag and labeled via the anti-GFP nanobody "enhancer".

(**E**) Representative smFRET trajectories observed for IL-7RΔECD WT and V253G, respectively. Scale bar: 5 μm.

(**F**) smFRET efficiency histograms for IL-7RΔECD WT and mutants with fit by multiple Gaussian functions (top), and localization map of co-localized donor and acceptor signals from 150 consecutive frames color-coded according to the corresponding FRET efficiency (bottom). Scale bar: 5 μm.

(**G**) Cell surface density of IL-7RΔECD WT and mutants observed in different smFRET dimerization experiments.

(**H**) Normalized smFRET trajectory length histograms for IL-7RΔECD WT and mutants.

(**I**) Comparison of mean dimer lifetime for IL-7RΔECD WT and mutants as quantified from smFRET trajectory length analysis.

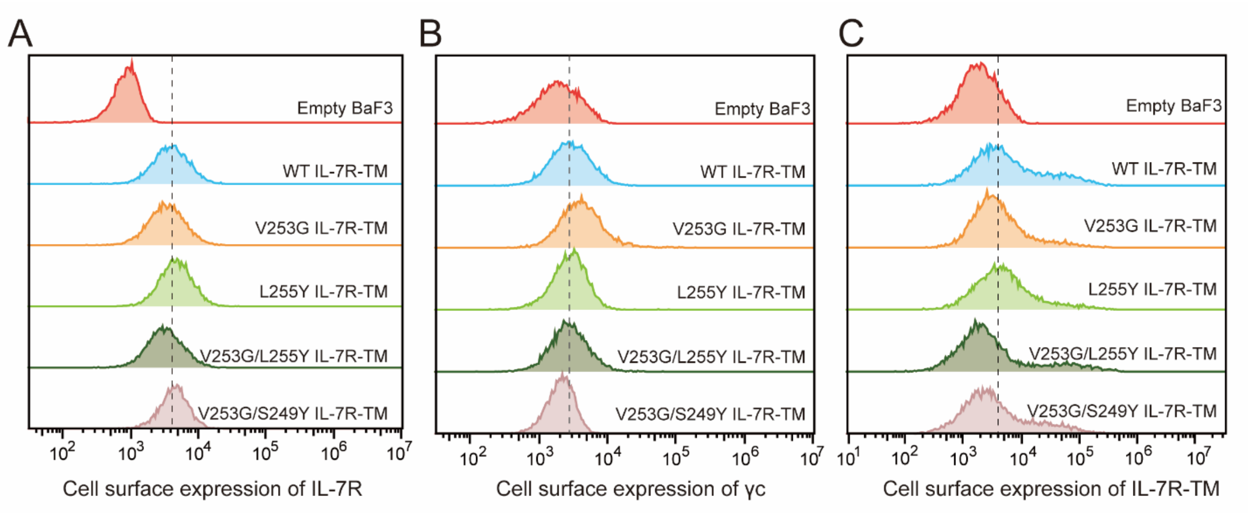


**Fig. S9. Flow cytometry analysis of cell surface expression of IL-7R, γc and different designed TMDs.**

(**A**) Flow cytometric analysis of IL-7R membrane expression in BaF3 cells co-expressing IL-7R and γc. Untransfected (Empty) BaF3 cells were used as negative controls.

(**B**) Flow cytometric analysis of γc membrane expression in BaF3 cells co-expressing IL-7R and γc. Untransfected (Empty) BaF3 cells were used as negative controls.

(**C**) Flow cytometric analysis of different designed IL-7R TMD membrane expression in BaF3 cells co-expressing V253G IL-7R and γc using APC-Flag antibody.

**Table S1. NMR structure calculation and refinement statistics**

| **NMR distance and dihedral constraints** [a] | |
|---|---|
| Distance constraints from NOE | 214 |
| Short-range ($\lvert i-j \rvert \leq 4$) | 96 × 2 |
| Long-range intramolecular ($\lvert i-j \rvert \geq 5$) | 0 |
| Intermolecular | 22 |
| Total dihedral angle restraints [b] | 100 |
| $\phi$ (TALOS) | 50 × 2 |
| $\psi$ (TALOS) | 50 × 2 |
| **Structure statistics** [c] | |
| Violations (mean ± s.d.) | |
| Distance constraints (Å) | 0.050 ± 0.007 |
| Dihedral angle constraints (º) | 0.135 ± 0.037 |
| Deviations from idealized geometry | |
| Bond lengths (Å) | 0.005 ± 0.000 |
| Bond angles (º) | 0.558 ± 0.013 |
| Impropers (º) | 0.317 ± 0.023 |
| Average pairwise r.m.s. deviation (Å) [d] | |
| Heavy | 1.246 |
| Backbone | 0.631 |

[a] The numbers of restraints for the IL-7Rtm-V253G are summed over residues 236-266.

[b] Backbone $\phi$ and $\psi$ restraints and their respective uncertainties were obtained from the "GOOD" dihedrals generated by the TALOS+ program [53] based on the backbone chemical shift values.

[c] Statistics are calculated and averaged over an ensemble of the 15 lowest energy structures out of 100 calculated structures.

[d] The precision of the atomic coordinates is defined as the average r.m.s. difference between the 15 final structures and their mean coordinates. The calculation only includes the ordered regions of the protein: residues 240-264 for IL-7Rtm-V253G.

**Table S2. Overview of MD Simulation Systems and Sampling Scheme**

| System name | Number of initial models | Number of replicates per initial model | Simulation time (ns) per replicate |
|---|---|---|---|
| **Extended MD simulations (Temperature = 310 K)** | | | |
| Mutant-G253 | 1 | 3 | 2000 |
| | | 7 | 1000 |
| WT-V253 | 1 | 3 | 2000 |
| | | 7 | 1000 |
| WT-L255 | 10[a] | 3 | 1000 |
| **Temperature Scan Simulations (Temperature = 310 K - 350 K)** [b] | | | |
| Mutant-G253 | 1 | 3 | 200 + 100 |
| WT-V253 | 1 | 3 | 200 + 100 |
| WT-L255 | 10[a] | 3 | 200 + 100 |

[a] Best scoring 10 Boltz-2 Models were considered.

[b] The temperature was linearly increased from 310 K to 350 K in 200 ns and kept at 350 K for an additional 100 ns.
